# Systemic Proteome Phenotypes Reveal Defective Metabolic Flexibility in Mecp2 Mutants

**DOI:** 10.1101/2023.04.03.535431

**Authors:** Stephanie A. Zlatic, Erica Werner, Veda Surapaneni, Chelsea E. Lee, Avanti Gokhale, Kaela Singleton, Duc Duong, Amanda Crocker, Karen Gentile, Frank Middleton, Joseph Martin Dalloul, William Li-Yun Liu, Anupam Patgiri, Daniel Tarquinio, Randall Carpenter, Victor Faundez

**Affiliations:** Departments of Cell Biology, Emory University, Atlanta, GA, USA, 30322; Biochemistry, Emory University, Atlanta, GA, USA, 30322; Program in Neuroscience, Middlebury College, Middlebury, Vermont 05753; Department of Neuroscience and Physiology, SUNY Upstate Medical University, Syracuse, NY 13210, USA; Pharmacology & Chemical Biology, Emory University, Atlanta, GA, USA, 30322; Center for Rare Neurological Diseases. Norcross, GA 30093, USA; Rett Syndrome Research Trust, Trumbull, CT 06611, USA

## Abstract

Genes mutated in monogenic neurodevelopmental disorders are broadly expressed. This observation supports the concept that monogenic neurodevelopmental disorders are systemic diseases that profoundly impact neurodevelopment. We tested the systemic disease model focusing on Rett syndrome, which is caused by mutations in *MECP2*. Transcriptomes and proteomes of organs and brain regions from *Mecp2*-null mice as well as diverse *MECP2*-null male and female human cells were assessed. Widespread changes in the steady-state transcriptome and proteome were identified in brain regions and organs of presymptomatic *Mecp2*-null male mice as well as mutant human cell lines. The extent of these transcriptome and proteome modifications was similar in cortex, liver, kidney, and skeletal muscle and more pronounced than in the hippocampus and striatum. In particular, *Mecp2*- and *MECP2*-sensitive proteomes were enriched in synaptic and metabolic annotated gene products, the latter encompassing lipid metabolism and mitochondrial pathways. *MECP2* mutations altered pyruvate-dependent mitochondrial respiration while maintaining the capacity to use glutamine as a mitochondrial carbon source. We conclude that mutations in *Mecp2*/*MECP2* perturb lipid and mitochondrial metabolism systemically limiting cellular flexibility to utilize mitochondrial fuels.

## Introduction

Rett syndrome is a neurodevelopmental disorder caused by loss-of-function mutations in the epigenetic transcriptional regulator *MECP2* (Amir et al., 1999; Chahrour and Zoghbi, 2007; Sandweiss et al., 2020). To date, the study of human and animal models of Rett and other neurodevelopmental disorders has primarily focused on the brain. The devastating nature of the neurological and behavioral phenotypes that characterize this and other monogenic neurodevelopmental disorders has rightly justified this focus on neuronal mechanisms of disease (Chahrour and Zoghbi, 2007). However, the protein products of the *MECP2* and other neurodevelopmental disease genes are widely expressed, and non-neuronal disease phenotypes (Schmidt et al., 2018), support the proposition that Rett and other neurodevelopmental disorders may be systemic diseases that profoundly impact neurodevelopment (Borloz et al., 2021; Kyle et al., 2018; Vashi and Justice, 2019). Such conceptualization could enhance biomarker and therapeutic discovery expanding the focus to organs, systems, and fluids beyond the brain and cerebrospinal fluid. Other tissues and cells from patients and animal models could act as a proxy for brain disease, thus empowering discovery of disease mechanism and therapies.

The systemic metabolic disease model of Rett syndrome was formulated before the identification of *MECP2* as the causative gene defect (Kyle et al., 2018). This idea originated from the Rett syndrome resemblance to mitochondrial diseases and patient alterations pyruvate and lactate levels in biofluids (Jellinger et al., 1988; Neul et al., 2020; Philippart, 1986). In fact, subjects with MECP2 mutations can manifest disease as a mitochondrial disorder (Kohda et al., 2016). Numerous studies in humans and in animal models of this disease document the systemic nature of Rett syndrome (Borloz et al., 2021; Kyle et al., 2018; Vashi and Justice, 2019). Humans affected by Rett syndrome have symptoms associated with the respiratory, cardiovascular, digestive, endocrine and musculoskeletal systems (Borloz et al., 2021; Kyle et al., 2018; Vashi and Justice, 2019). *Mecp2* mutant mouse models identify phenotypes in adipose tissue, respiratory, and digestive organs that are cell-autonomous and cannot be attributed simply to defects in organ innervation. Cell-type specific ablation of the *Mecp2* gene in adipocytes, lung epithelial cells, and hepatocytes alters cellular metabolism at the expense of cholesterol, fatty acid, and mitochondrial pathways (Kyle et al., 2016; Liu et al., 2020; Vashi et al., 2021). These *Mecp2* mutant phenotypes can be corrected with genetic or pharmacologic therapeutics that target specific pathways (Buchovecky et al., 2013; Enikanolaiye et al., 2020). This growing evidence supports the hypothesis that Rett syndrome is a systemic metabolic disease (Borloz et al., 2021; Kyle et al., 2018; Vashi and Justice, 2019).

We unbiasedly evaluated the systemic effects of *Mecp2* mutations by defining the transcriptome and proteome of organs and brain regions from *Mecp2*-null mice. Profound alterations of transcriptomes and proteomes were identified in the brain and organs of *Mecp2^tm1.1Bird/y^* animals before the appearance of overt phenotypes. Alterations in the transcriptome and proteome are most pronounced in the brain cortex, liver, kidney and skeletal muscle. *Mecp2*-sensitive proteomes are enriched in metabolic annotated gene products for lipid metabolism and mitochondrial pathways across diverse tissues. Importantly, the proteome of the cortex, the most affected brain region, was comparably enriched for both synaptic and mitochondrial annotated proteins. These analyses indicated that a mitochondrial and lipid metabolic hub is perturbed in *Mecp2* tissues and impairs pyruvate-dependent mitochondrial function. We conclude that mutations in *Mecp2*/*MECP2* systemically alters lipid and mitochondrial metabolism. We propose that compromised metabolic flexibility, flexibility in the utilization of carbon sources by mitochondria, is a systemic mechanism of neurodevelopmental disease.

## Results

### Organ-Specific Divergence of the *Mecp2* Mutant Transcriptomes and Proteomes

Bulk transcriptome and proteome analyses of microdissected brain regions (cortex, hippocampus, striatum), liver, kidney, and skeletal muscle from wild type and *Mecp2^tm1.1Bird/y^* mice were analyzed to define systemic molecular phenotypes in *Mecp2*-null mice. To define molecular changes preceding overt disease, mutant mice were sacrificed at 45 days of age. Approximately 80% of *Mecp2^tm1.1Bird/y^* mice survive to this age, fifty percent of the mice are neurologically symptom-free, and there is no overt organ pathology, such as fatty liver (Guy et al., 2001).

The *Mecp2*-null mutation affected the transcriptome of all tissues analyzed (Fig. 1A-B). The top-ten most increased or decreased transcripts differed across tissues (Fig. 1B). Whole transcriptomes segregated organs and brain regions by genotype as revealed by Uniform Manifold Approximation and Projection analysis (Fig. 1C, UMAP). Each one of the *Mecp2^tm1.1Bird/y^* tissues analyzed had a similar percentage of transcripts with either increased or decreased steady-state levels (Fig. 1A, red and blue symbols respectively). However, the total number and identity of differentially expressed transcripts varied among these tissues (Fig. 1A-B). The most affected tissues were liver and kidney, with ∼7-8 % of all quantified mRNAs with steady-state levels modified by the *Mecp2* mutation (Fig. 1A). In contrast, microdissected brain regions were the least affected tissues, with altered expression of 1.4-4.4% of the mRNAs (Fig. 1A). Transcriptome similarity matrices comparing across brain regions and genotypes showed that the *Mecp2^tm1.1Bird/y^* mutation caused the most pronounced differences between genotypes in cortex as compared to hippocampus and striatum (Fig. 1D). We chose the *Mecp2^tm1.1Bird/y^* cortical transcriptome to determine its replicability. The Pearson correlation among *Mecp2^tm1.1Bird/y^* differentially expressed mRNAs was 0.68 in two independent cohorts of microdissected cortices and transcriptional analyses (Fig. 1E). A similar correlation was obtained comparing our findings with either public whole-cell or nuclear cortical transcriptome datasets obtained in the same *Mecp2^tm1.1Bird/y^* mouse at a similar age (Fig. 1F and G, Pearson=0.69 and 0.59) (Clemens et al., 2020). We previously validated mRNA steady-state differences across wild type brain regions against Allen mouse brain transcriptome data (r=0.69) (Wynne et al., 2021). These multipronged validation approaches show that our bulk transcriptome data fulfill reproducibility standards.

**Figure 1.**
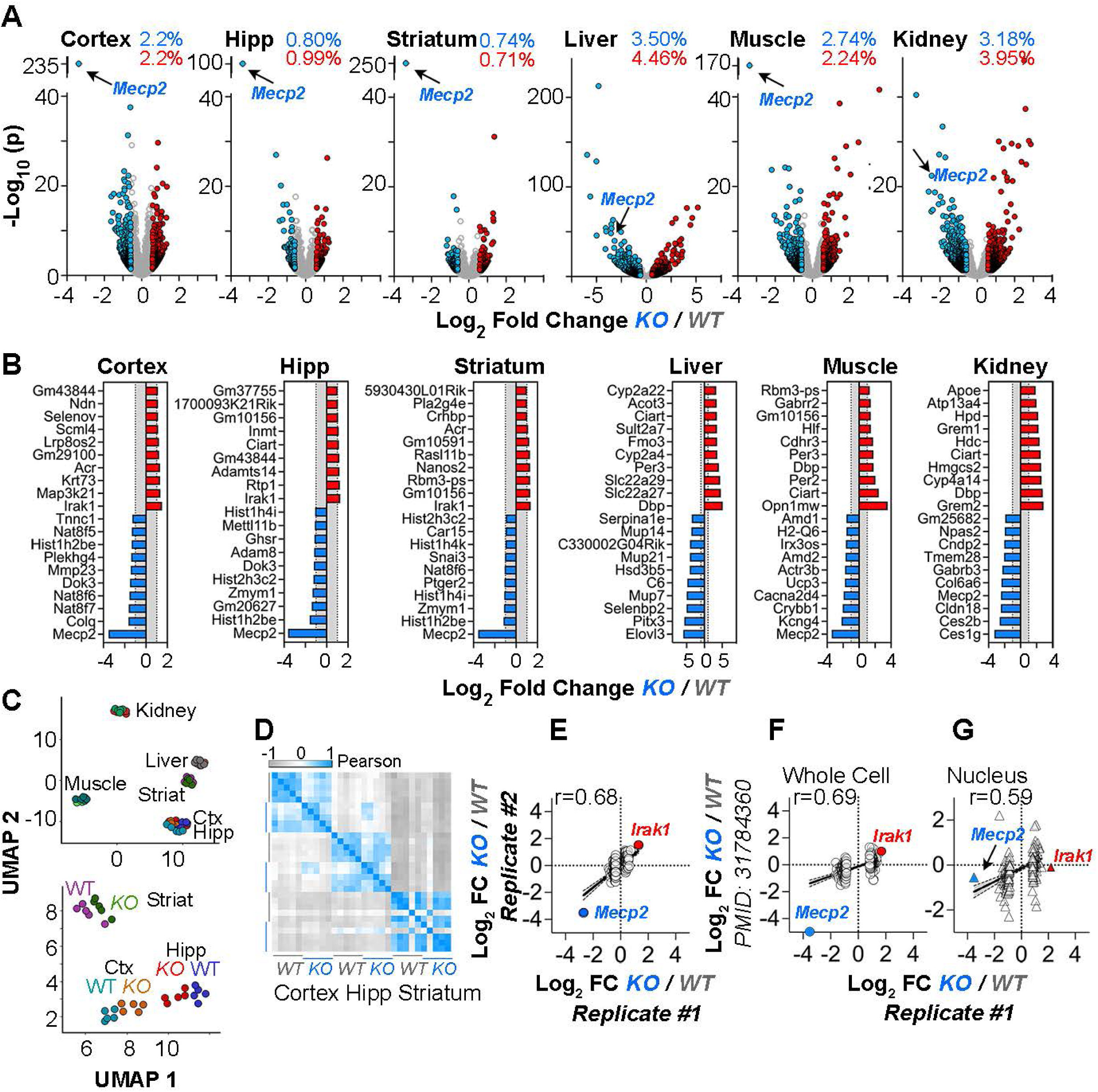
The *Mecp2^tm1.1Bird/y^* Tissue-Specific Transcriptomes. **A.** Volcano plots of microdissected cortex, hippocampus, striatum, liver, skeletal muscle, and kidney. Fold of change threshold is 1.5-fold and corrected p value cut-off is set at 0.05. Red and blue symbols represent transcripts whose steady-state levels are increased and decreased in mutants, respectively (n=5 animals of each genotype). **B**. Top 10 most increased and decreased transcripts in *Mecp2* null tissues. Data expressed as log2 fold of change. Gray area depicts the −1 to +1 log2 interval. **C.** UMAP analysis wild type and mutant tissue whole transcriptomes. **D**. Pearson correlation similarity matrix of wild type and mutant microdissected brain regions whole transcriptomes. **E**. Pearson correlation of two independent microdissected cortical transcriptome datasets (n=5 animals of each genotype per dataset). **F-G.** Pearson correlation of microdissected cortical transcriptome Replicate #1 with bulk cortical tissue or nuclear fraction transcriptomes reported by Clemens et al. (Clemens et al., 2020). This dataset was chosen because of the age, the use of the *Mecp2^tm1.1Bird/y^* mouse strain, and brain region analyzed match those used by us. The top 100 most increased and decreased transcripts in the Replicate #1 mutant dataset were compared with Replicate #2 in E and with Clemens et al datasets in F-G. The two most changed transcripts *Mecp2* and *Irak1* are antipodes shared by all datasets.

Since the *Mecp2* mutation alters the transcriptome across all organs, pathway analysis were performed using whole tissue transcriptomes and the Gene Set Enrichment Analysis (GSEA) tool (Subramanian et al., 2005) to identify processes/mechanisms commonly affected across the different tissues. These analyses did not identify significant pathways shared across mutant tissue transcriptomes (data not shown). We therefore focused on *Mecp2*-sensitive mRNAs shared by all tissues to define common pathway phenotypes. The expression of 20 mRNAs was altered by the *Mecp2* mutation in all tissues (Fig. S1A). The overlap among brain regions was also discrete with 69 differentially expressed mRNAs whose levels were modified in all three brain regions. Of these 69 overlapping brain mRNAs, 38 were identified as congruently changed in independent cortical datasets listed above (Fig. S1B, bold red and blue fonts). These 69 differentially expressed mRNAs were not enriched in significant ontologies or pathways. Thus, *Mecp2* mutations alter the transcriptome systemically with minimal transcript overlap across tissues, and no apparent pathway enrichment in brain or other tissues.

The performance of the transcriptome to identify systemic pathways sensitive to *Mecp2* gene defects prompted us to ask whether the proteome could better predict pathways affected by *Mecp2* mutations. The proteome was assessed using Tandem Mass Tagging (TMT) quantitative mass spectrometry of microdissected brain regions, liver, and skeletal muscle to measure the extent of the *Mecp2*-sensitive proteome (Fig. 2A) (Thompson et al., 2003). A total of 10,087 proteins were simultaneously quantified across these tissues in both genotypes; corresponding to 5,332 in cortex, 5,540 in hippocampus, 2,874 in striatum, 8,579 in liver, and 5,842 proteins in skeletal muscle. In distinct contrast to the transcriptome, the cortex proteome was the most affected tissue with the steady-state levels of 322 proteins, 6% of the quantified cortex proteome, altered in *Mecp2* mutants (Fig. 2A-B). Liver and skeletal muscle proteomes were moderately affected with 130 (1.5%) and 167 (3.2%) protein levels altered, respectively (Fig. 2A-B). In contrast, the proteomes of hippocampi and striata were minimally disrupted by the *Mecp2* mutation (Fig. 2A-B). Divergences between the tissue transcriptomes and proteomes prompted us to assess the similarity of the transcriptome and proteome across tissues (Fig. 2C). Steady-state level differences from all significantly changed *Mecp2*-sensitive transcripts were compared to the corresponding protein levels. Although the proteome and transcriptome correlated in wild type cortex (Fig. 2D, r=0.38), the *Mecp2*-sensitive cortical transcriptome-proteome (encompassing 1532 mRNA-protein pair quantifications) did not correlate (Fig. 2C, r=0.0058). The lack of correlation in mutant cortex is in stark contrast with the correlation between the liver *Mecp2*-sensitive transcriptome and proteome, where the Pearson correlation was 0.69 for a large size set of transcript-protein pairs (Fig. 2C, n=767 pairs), a value comparable to the correlation between the liver proteome and transcriptome in wild type tissue (Fig. 2D, r=0.50). We hypothesized that differences in the expression of small regulatory microRNAs could account for divergent correlations between the proteome and the transcriptome in cortex and liver of *Mecp2^tm1.1Bird/y^* animals (Fig. S2). However, the number of microRNAs differentially expressed in these *Mecp2* mutant tissues and their predicted target mRNAs is unlikely to account for the poor correlation of the transcriptome and proteome in *Mecp2* mutant cortex. There were few exceptions, such as Slc25a51, a mitochondrial nicotinamide adenine dinucleotide (NAD^+^) transporter (Luongo et al., 2020), which is a predicted target of miR-182-5p, a microRNA increased in the microdissected cortex of mutant mice (Fig. S2). These results demonstrate that factors other than the transcriptome specify the steady-state proteome in *Mecp2^tm1.1Bird/y^* brain.

**Figure 2.**
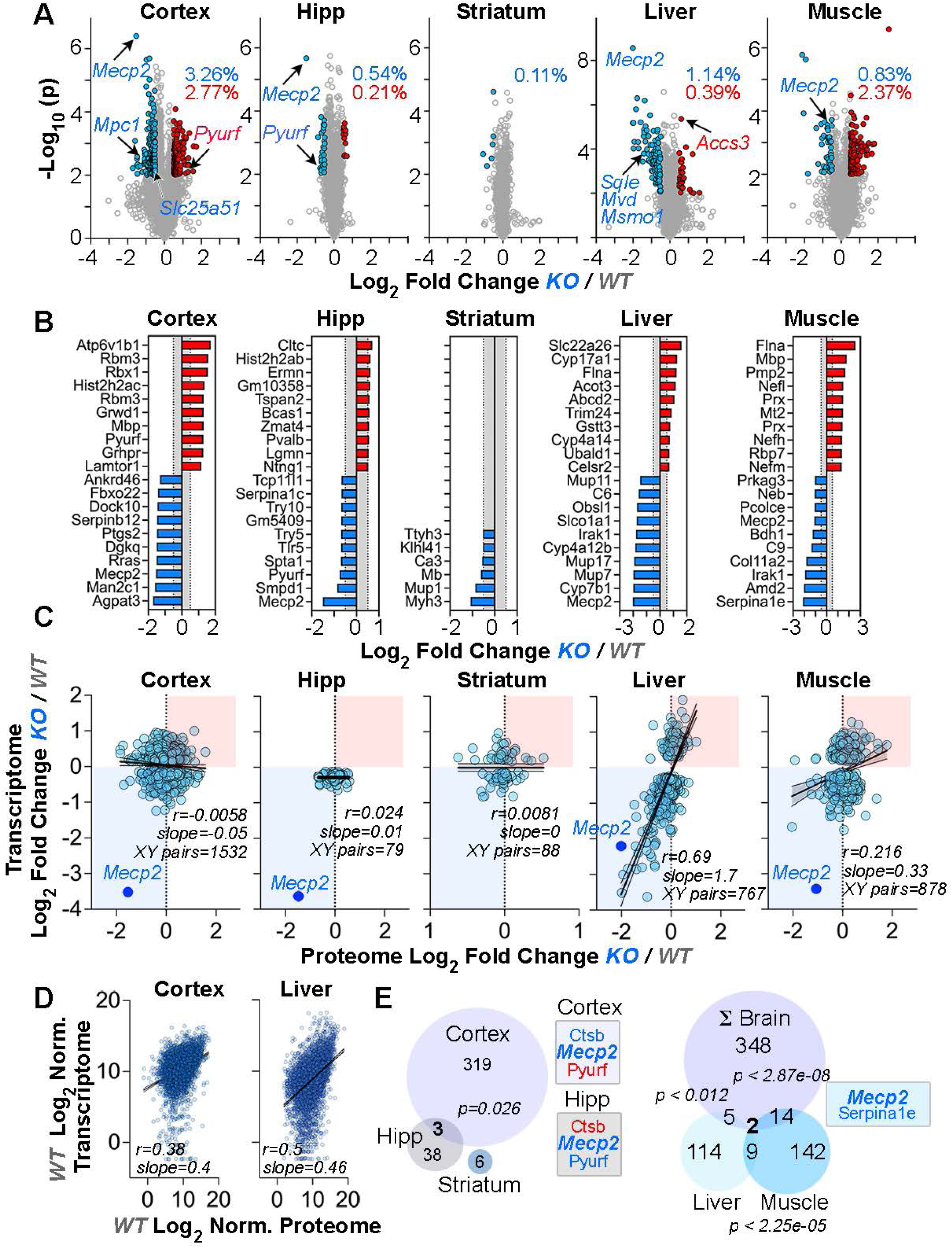
The *Mecp2^tm1.1Bird/y^* Tissue-Specific Proteomes. **A.** Volcano plots of microdissected cortex, hippocampus, striatum, liver, skeletal and muscle. Fold of change threshold is 1.5-fold and p value cut-off is set at 0.05. Red and blue symbols represent proteins whose steady-state levels are increased and decreased in mutants, respectively (n=5 animals of each genotype). **B**. Top 10 most increased and decreased proteins in *Mecp2* null tissues. Data expressed as log2 fold of change. Gray area depicts the −0.5 to +0.5 log2 interval. **C.** Pearson correlation of differentially expressed transcripts thresholded just by corrected p values <0.05 and their matching protein pair for each tissue. **D**. Spearman correlation of the whole quantified proteome and transcriptome for microdissected cortex and liver. **E**. Venn diagrams of overlapping proteins with altered steady-state levels in mutant brain regions and the sum of all differentially expressed proteins in brain (ΣBrain) compared to muscle and liver mutant proteomes. Hypergeometric p values for the different datasets overlaps. See Supplementary Table 2.

In order to identify universal proteome phenotypes, we focused on proteins that were altered across multiple *Mecp2* mutant tissues. There was minimal overlap among differentially expressed proteins across *Mecp2* mutant brain regions (Fig. 2A-B, E, and S3A), as well as between brain regions and other tissues (Fig. 2B, E, and S3A). For example, the mutant cortex and hippocampus shared Mecp2 and two other proteins, cathepsin B (Ctsb) and the mitochondrial complex I assembly factor PreY (Pyurf) (Moutaoufik et al., 2019; Rensvold et al., 2022). However, levels of these two proteins were affected in opposite directions in microdissected cortex as compared to hippocampus (Fig. 2A-B and E). Other than Mecp2, a single protein was decreased in all tissues the alpha-1-antitrypsin isoform Serpina1e (Fig. 2E and S3A). Serpina1e was in lowest abundance quartile of the *Mecp2*-sensitive cortex proteins while it was in the uppermost quartile in liver and muscle (Fig. S3B). Of note, Serpina1e protein levels were previously reported to be reduced levels in the CSF of *Mecp2* mutant mice (Zlatic et al., 2022). Genetic defects in *Mecp2* cause widespread non-overlapping modifications of the transcriptome and proteome across tissues. Furthermore, the levels of a *Mecp2*-sensitive mRNA variably correlate with the corresponding protein level in tissues with the exception of nervous tissue, where there was no correlation.

### *Mecp2/MECP2* Mutant Transcriptomes and Proteomes Enrich Metabolic Ontologies Across Organs and Cells

Despite the lack of correlation across tissue transcriptomes, we reasoned that combining *Mecp2* mutant tissue proteomes and transcriptomes may enable identification of shared pathways. *Mecp2*-sensitive proteomes were analyzed with the ClueGo tool and transcriptomes and proteomes were interrogated for pathway convergence with the Metascape tool (Bindea et al., 2009; Zhou et al., 2019). Metascape analysis of parental ontologies for tissue transcriptomes and proteomes identified shared ontologies for metabolic process (GO:0008152, Fig. S4A) and response to stimulus (GO:0050896, Fig. S4A). These parental ontology nodes gave rise to descendant terms with different degrees of representation across tissues and ‘omic’ datasets (Fig. S4B). For example, organic hydroxy compound metabolism was represented in all but one of the tissue omics and this term co-clustered with other metabolic ontologies including small molecule catabolic process (GO:0044282 -log_10_ p value 31.963) and metabolism of lipids (Fig. S4B, R-HSA-556833 -log_10_ p value 55.139). A similar outcome was obtained by interrogating tissue proteomes with the ClueGO tool (Fig. S4C). The strongest ontologies were cell responses to stimuli, synaptic, lipid metabolism, mitochondria, and peroxisomes (Fig. S4C). These ontologies were represented in all tissue proteomes to a different extent (Fig. S4D). Lipid metabolism ontologies preferentially enriched in liver while synaptic and mitochondrial ontologies were enriched in brain (Fig. S4C-D). In fact, we found a decreased steady-state level of 10 enzymes required for cholesterol synthesis in *Mecp2*-null liver (Fig. S5). Brain and liver proteomes also shared large numbers of synapse-annotated (SynGO), lipid (GO:0006629), and mitochondrial (Mitocarta 3.0) annotated proteins (Fig. S4E) (Koopmans et al., 2019; Rath et al., 2021). The degree of enrichment in synaptic and mitochondrial annotated proteins in *Mecp2* mutant brains was similarly significant with a 2.4-2.8-fold enrichment (Fig. S4F, p<2E-8, Exact hypergeometric probability). Similar results were identified analyzing a published *Mecp2* mutant proteome obtained from the whole isocortex of the alternate null allele (*Mecp2 ^Jae/y^*) at a later, symptomatic, age (P60, Fig. S6A-B) (Pacheco et al., 2017).

We next asked whether metabolic ontologies found in mouse *Mecp2* mutant tissue proteomes were a universal feature of *MECP2* mutants irrespective of the biological systems scrutinized. We performed TMT quantitative mass spectrometry across human *MECP2*-null cells and their isogenic controls (Fig. 3). The variability between biological replicates from wild type and *MECP2*-null iPSC-derived human neurons was too large to permit identification of *MECP2*-sensitive proteomes in these cells (unpublished data). We therefore analyzed three alternative human cell lines; the chronic myelogenous leukemia male cell line HAP1; and two neuronal female cell lines, LUHMES postmitotic neurons (LUHMES, Lund human mesencephalic neurons), and the SH-SY5Y neuroblastoma cells (Biedler et al., 1978; Kotecki et al., 1999; Scholz et al., 2011). Cells of the myeloid lineage express MECP2 protein at levels comparable to brain and show alterations of the transcriptome in Rett patients (Pecorelli et al., 2013; Samaras et al., 2020; Schmidt et al., 2018). We quantified 5958 proteins in LUHMES, 8086 proteins in SH-SY5Y, and 6463 proteins in HAP1 cells. These proteomes identified 186, 323, and 106 proteins whose steady-state levels were altered in *MECP2*-null HAP1 (male), LUHMES (female), and SH-SY5Y cells (female), respectively (Fig. 3A-B). Metascape analysis of parental ontologies shared across cell lines and organ proteomes indicated that the strongest shared ontology was metabolic process (Fig. 3C, GO:0008152, -log_10_ p value 13, 38.1, and 29.5; ΣBrain, HAP1 and liver, respectively). This parental term enclosed multiple mitochondrial and lipid metabolism ontologies. Among these, the two most widely represented ontologies across *Mecp2*/*MECP2* proteomes were generation of precursor metabolites and energy with 50% mitochondrial annotated proteins (Fig. 3D, GO:0006091 -log_10_ p value 13, 17.9, and 2.3; ΣBrain, HAP1 and liver, respectively) and nucleobase-containing small molecule metabolic process with 35% mitochondrial annotated proteins (Fig. 3D, GO:0055086 -log_10_ p value 10, 3.6, and 6.5; ΣBrain, HAP1 and liver, respectively). The HAP1 and mouse brain proteomes were the datasets with the highest number mitochondrial annotated proteins (Mitocarta 3.0, Fig. S7A) (Rath et al., 2021). We measured the degree of overlap between human cell line and mouse brain mutant proteomes (Fig. S7B, Enrichment) and determined that the *MECP2* HAP1 proteome had the highest overlap normalized by dataset size with either the *Mecp2* mouse brain proteome or the Mitocarta 3.0 proteome among the three human cell lines tested. The *MECP2* mutant HAP1 proteome was enriched in *Mecp2* sensitive mouse brain proteins 4.3-times above what was expected by chance (Fig. S7B, p<5E-4, Exact hypergeometric probability). These results demonstrate that metabolic and mitochondrial ontologies are a common phenotype across diverse *Mecp2*/*MECP2* deficient biological systems. These results suggest that mitochondrial and metabolic processes are preponderantly disrupted by genetic defects in *MECP2*/*Mecp2*.

**Figure 3.**
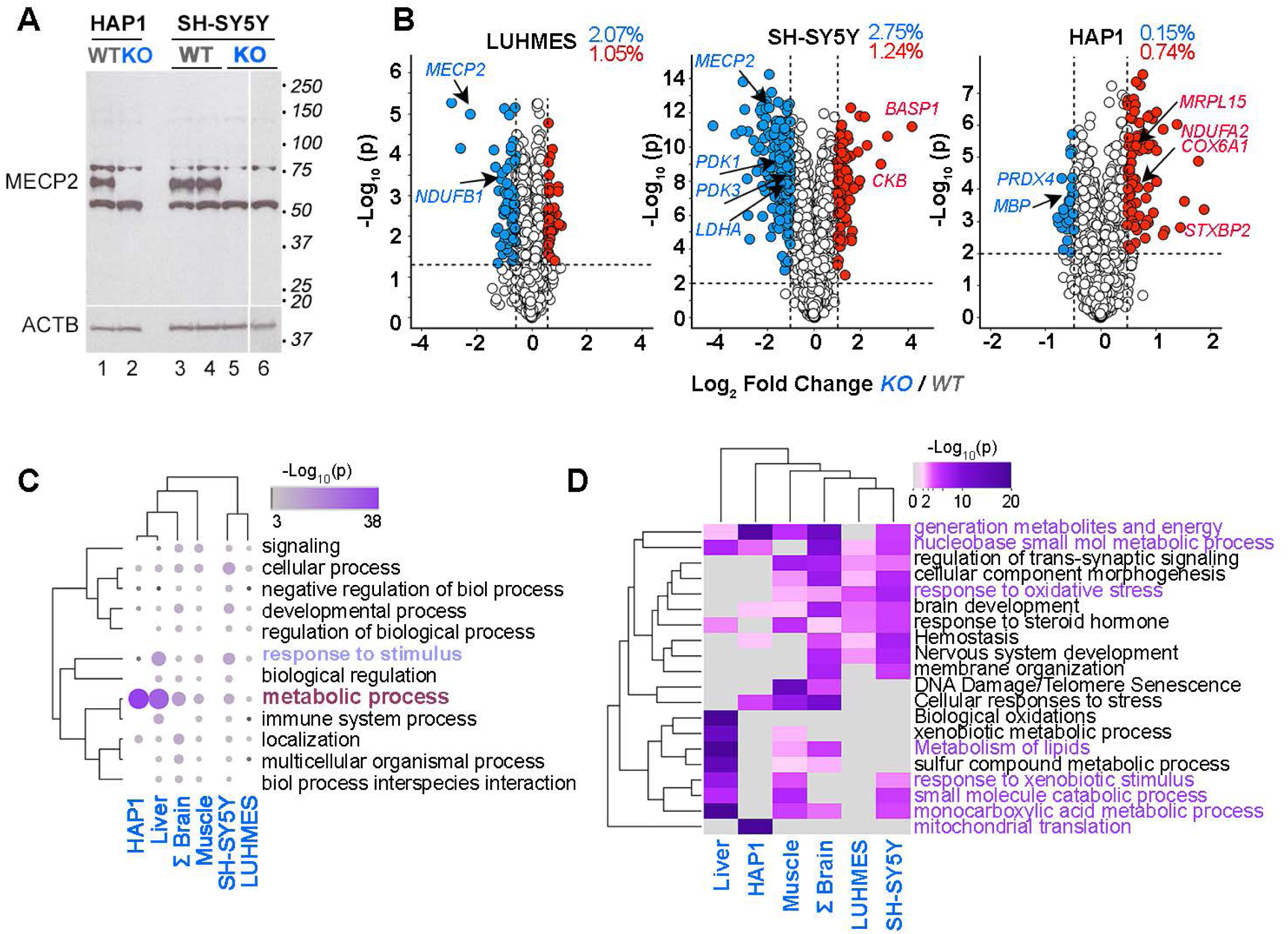
Human Cell Lines *MECP2* Proteomes Converge on Metabolic and Mitochondrial Ontologies. **A**. Immunoblots of HAP1 and SH-SY5Y cell lines control and *MECP2* edited. Actin B (ACTB) was used as loading control. Lanes 3-4 and 5-6 represent two independent wild type and null cell lines respectively. **B.** Volcano plots of LUHMES, SH-SY5Y, and HAP1 wild type and *MECP2*-null cells. Fold of change threshold is 1.5-2-fold and corrected p value cut-off is set at 0.05 for LUHMEs cells, and 0.001 for HAP1 and SH-SY5Y cells. Red and blue symbols represent proteins whose steady-state levels are increased and decreased in mutants, respectively (n=5 independent replicates per genotype, except for LUHMEs cells n=3). **C.** Parental GO term ontologies represented in the differentially expressed proteomes of mutant mouse tissues and human cell lines. Clustering performed with Pearson correlation. ΣBrain represents the added microdissected cortex and hippocampal *Mecp2* proteomes. **D**. Ontologies, KEGG terms, and pathways enriched and shared by datasets used in C. Accumulative hypergeometric p-values and enrichment factors were calculated and used for filtering. Significant terms were clustered into a tree using kappa-statistical similarities among their gene memberships. Grey denotes no representation of the dataset in an ontology.

### *Mecp2/MECP2* Mutations Differentially Disrupt Mitochondrial Proteins

Mitochondrial annotated proteins intersect synaptic, peroxisome, and lipid metabolism ontologies and processes. Mitochondria occupy 30% of the synapse volume and are necessary for synaptic transmission as well as for cholesterol and fatty acid metabolism (Chandel, 2021; Cheng et al., 2022; Martinez-Reyes and Chandel, 2020; Rangaraju et al., 2014; Wilhelm et al., 2014). Levels of 45 proteins (annotated to Mitocarta 3.0) changed only in cortex including the pyruvate carrier Mpc1, the NAD^+^ carrier Slc25a51, and Pyurf (Fig. S7C-D) (Rath et al., 2021). Coessentiality network analysis with the Fireworks tool (Fig. S7D) (Amici et al., 2021) revealed that the most altered mitochondrial proteins in *Mecp2* mutant cortex, such as Mpc1, Slc25a51, and Pyurf, shared functional relationships with respiratory chain complex I subunits (Fig. S7D Pyurf and Ndufab9 nodes) and proteins involved in pyruvate metabolism (Fig. S7D Mpc1 node). These results suggest a defect in the mitochondrial pyruvate metabolism.

Mpc1 is necessary for feeding pyruvate into the Krebs cycle, pyruvate incorporation into lipids, for maintaining steady-state levels of fatty acids and cholesterol in cells, and for normal synaptic transmission (Bowman et al., 2016; De La Rossa et al., 2022; Grenell et al., 2019). As such, the Mpc1 pyruvate transporter participates in processes and pathways uncovered by the *Mecp2* mutant proteome pathways and ontologies (Figs. 3C-D and S4C-D). Mpc1 levels were measured by immunoblot in *Mecp2^tm1.1Bird/y^* tissues as well as the three human *MECP2* mutant cell lines that we characterized by proteomics (Fig. 4). Mpc1 levels were either decreased or normal in microdissected cortex and hippocampus, respectively (Figs. S7C and 4A-B), thus confirming our mass spectrometry findings (Fig. 2A-B and S7C). Whether changes to levels of Mpc1 in micropunched-dissected tissue are recapitulated in whole isocortex extracts was assessed by immunoblot analysis of whole mouse isocortex extracts (Fig. 4C). The increased levels of Mpc1 in the whole cortex analyses suggests there is heterogeneity among regions of the isocortex (Fig. 4C). Mpc1 levels were increased in liver while Mpc1 and Mpc2 levels were decreased in mutant striated muscle (Fig. 4D-E). These results indicate that steady-state levels of pyruvate carriers are differentially affected in different organs and regionally within the brain.

**Figure 4.**
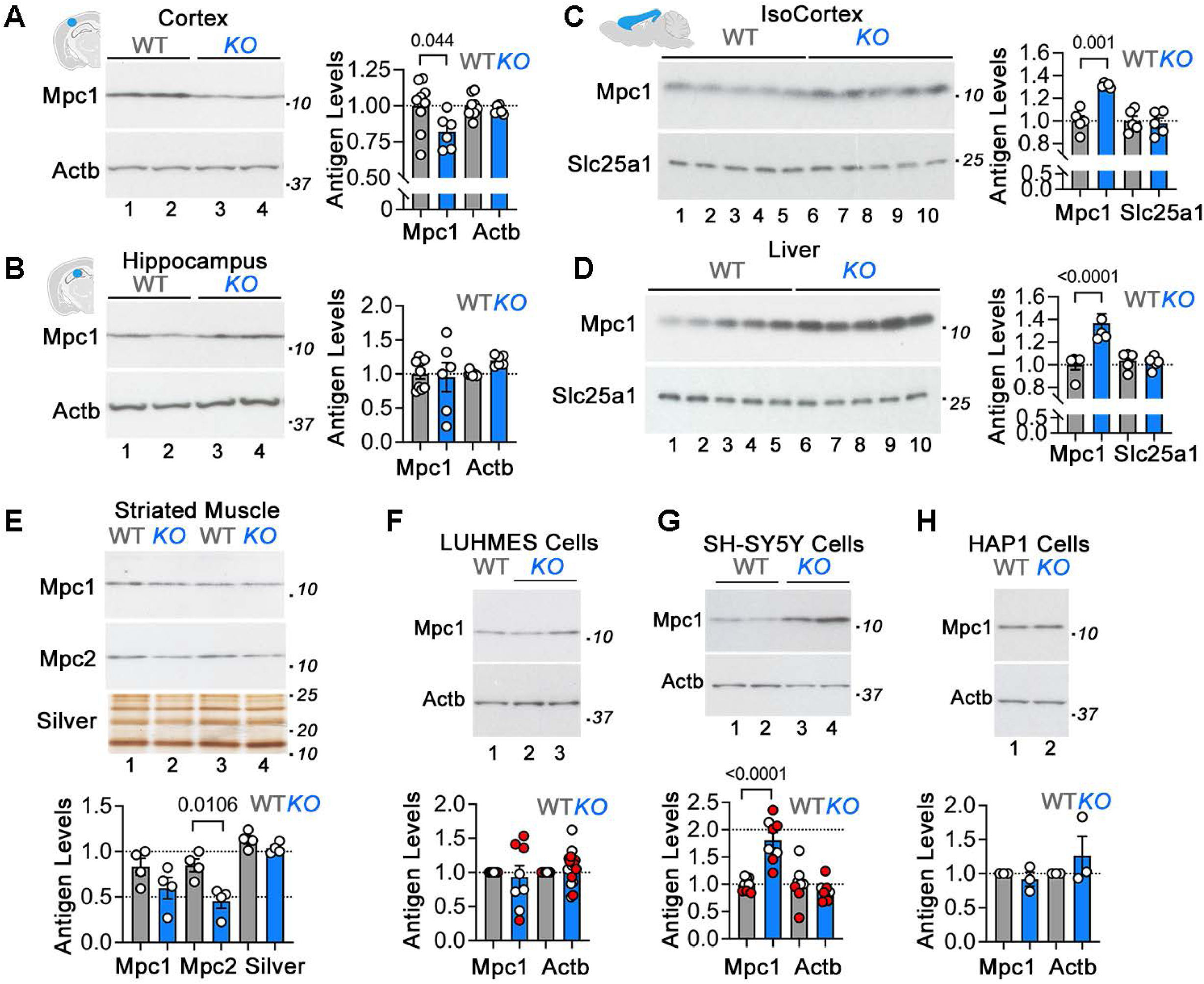
Alterations of Mitochondrial Pyruvate Carrier Levels in *Mecp2^tm1.1Bird/y^* Tissues and *MECP2* Mutant Cells. **A** to **H** show immunoblots and quantifications of the mitochondrial pyruvate transporters Mpc1 and Mpc2 in *Mecp2^tm1.1Bird/y^* tissue extracts (A, B, and E), *Mecp2^tm1.1Bird/y^* total membrane fractions (C and D), and MECP2-null human cell lines (F-H). Loading controls used were beta actin (Actb), the mitochondrial citrate transporter (Slc25a1), or silver stained SDS-PAGE. Quantifications are normalized to wild type antigen levels. Each dot represents an animal or a biological replicate. Red and white dots in F and G depict two independent *MECP2*-null clones. p values were obtained with two-sided permutation t-test with 5000 bootstrap samples. See Supplementary Table 2.

We then tested whether the variable levels of Mpc1 in mouse mutant tissues could be replicated by *MECP2*-null mutations across different human cell lines (Fig. 4F-H). The expression of Mpc1 in either MECP2 knockout differentiated LUHMES or HAP1 cells was not altered (Fig. 4F and H). In contrast, differentiated SH-SY5Y cells lacking MECP2 had 2-fold increase in Mpc1 expression (Fig. 4G). These finding indicate that *MECP2* mutations alter the expression of mitochondrial proteins and Mpc1 in a tissue and cell type-dependent manner. These findings suggest that *Mecp2*/*MECP2* mutant mitochondrial phenotypes such as respiration and pyruvate consumption may vary across cells and tissues.

### Mitochondrial Respiration is Altered in Cells Lacking MECP2

Tissue/cell type-specific modifications of the mitochondrial proteome in *Mecp2*/*MECP2*-null tissues and cell lines raise questions about mitochondrial functional phenotypes and the uniformity of these phenotypes across cell types. Mitochondrial respiration was measured in intact cells using Seahorse respirometry in complete media with glucose, pyruvate, and glutamine as carbon sources (Divakaruni et al., 2014). We postulated respiration phenotypes could be stunted by the anatomical heterogeneity inherent to primary cultures generated from whole isocortices as seen in the differing levels of Mpc1 between microdissected cortex and isocortex samples (compare Fig. 4A and C). Nevertheless, a discrete but significant decrease in the basal respiratory capacity was observed in primary neuronal cultures prepared from whole *Mecp2^tm1.1Bird/y^* and mosaic female *Mecp2^tm1.1Bird/+^* isocortices (Fig. 5A-B). There were no differences in the maximal respiration induced by uncoupling mitochondria with the protonophore carbonyl cyanide 4-(trifluoromethoxy)phenylhydrazone (FCCP. Fig. 5A and C). Differentiated mutant LUHMES cells (Fig. 5D-F) and HAP1 cells (Fig. 5J-L) had decreased basal and maximal mitochondrial respiration parameters. In contrast, a two-fold increase in both respiration parameters was observed in *MECP2* mutant SH-SY5Y cells (Fig. 5G-I). The same oxygen consumption phenotypes were observed when respiration was monitored continuously for 48 hours with Resipher platinum sensing probes in HAP1 and SH-SY5Y (Fig. S8A and C) (Grist et al., 2010; Wit et al., 2023). Neither we observed period oscillations in respiration in wild type and *MECP2* mutant cells (Fig. S8A and C), nor respiration phenotypes could be attributed to differences in cell numbers (Fig. S8A-B and C). These results support cell-type dependent mitochondrial respiration phenotypes in *Mecp2*/*MECP2* mutants.

**Figure 5.**
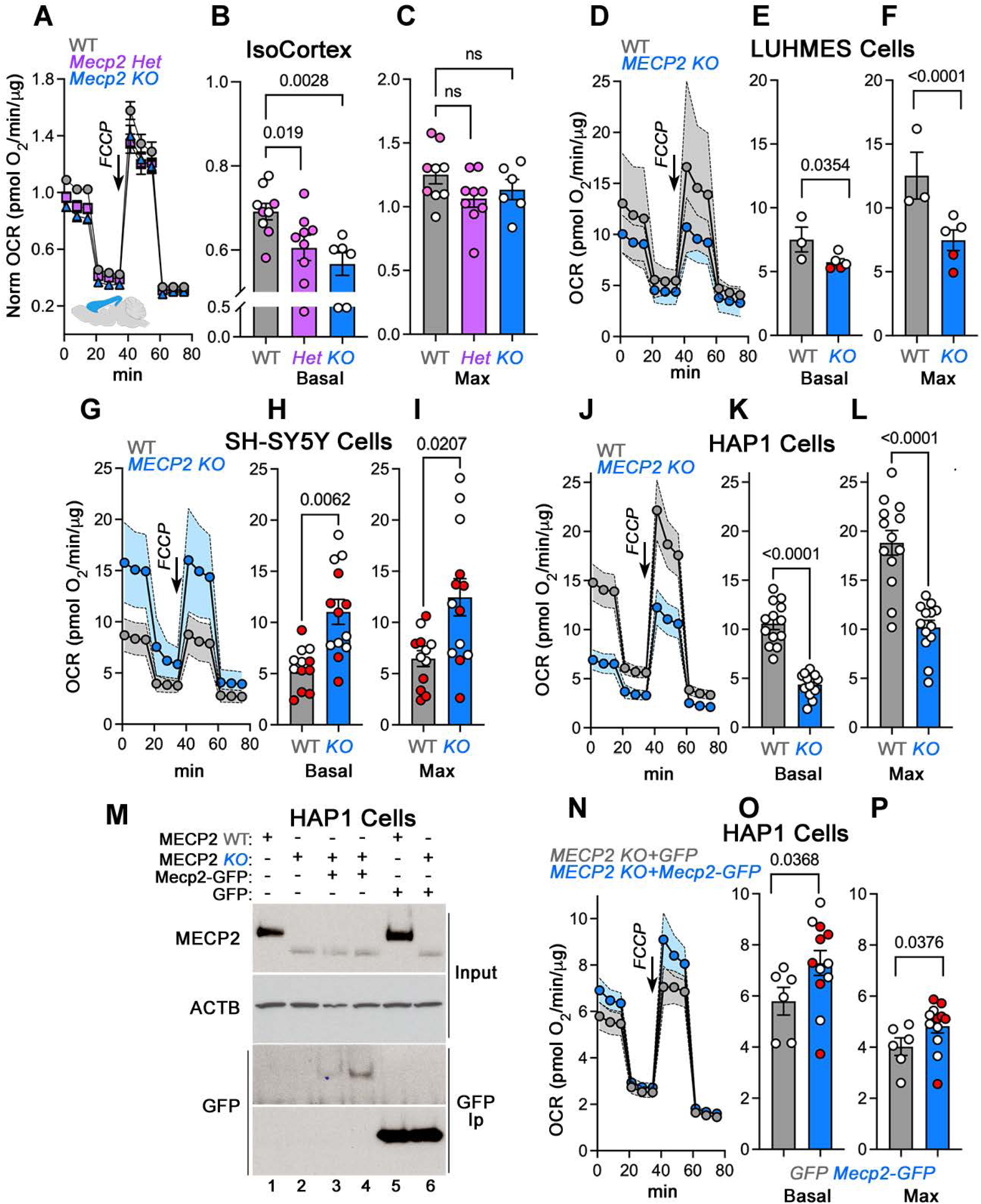
Mitochondrial Respiration is Altered in *Mecp2^tm1.1Bird/y^* Isocortex Primary Cultures and *MECP2* Mutant Cells. **A-C.** Primary cultured isocortex neurons from postnatal 1 cultured for 14 days were subjected to a Seahorse stress test in the presence of complete media. Oxygen consumption rate was normalized to wild type animal values at measurement 3. **B.** Depicts basal respiration and **C** presents maximal respiration elicited after FCCP addition (arrow). Each symbol corresponds to an independent animal, pink symbols mark female animals**. D** to **F** show mitochondrial stress test, basal and maximal respiration in wild type and *MECP2*-null differentiated LUHMES cells. **G** to **I** depict mitochondrial stress test, basal, and maximal respiration in wild type and *MECP2*-null differentiated SH-SY5Y neuroblastoma cells**. J-L** mitochondrial stress test, basal, and maximal respiration in wild type and *MECP2*-null HAP1 cells. **M**. Wild type (lane 1), stably expressing GFP (lane 5) and MECP2-null HAP1 cells (lane 2), stably expressing GFP (lane 6) or mouse *Mecp2*-GFP (lanes 3-4) were immunoprecipitated with anti-GFP antibodies and analyzed by immunoblot with antibodies against GFP. **N-P** depict mitochondrial stress test, basal, and maximal respiration in *MECP2*-null HAP1 cells stably expressing GFP (gray symbols) or *Mecp2*-GFP (blue symbols). **D, G, J** and **N** brackets and shaded area show average ± 95 confidence interval. Inflection after datapoint 3 marks addition of oligomycin and drop after datapoint 9 marks addition of rotenone and antimycin. All other data show average ± SEM. Each symbol is an independent biological replicate. Red symbols show an independent clone isolate. p values were obtained with two-sided permutation t-test with 5000 bootstrap samples.

We previously demonstrated that decreased mitochondrial respiration in mutant LUHMES cells can be restored by reintroduction of the *MECP2* gene (Zlatic et al., 2022). Similar gene re-expression experiments were performed in HAP1 cells. We chose to conduct these experiments in HAP1 cells because respirometry has the lowest sampling variability as assessed by 95% confidence intervals (compare shaded brackets in Fig. 5D, G and J). HAP1 MECP2 knock-out cells stably expressing mouse Mecp2-GFP (Fig. 5M, lanes 3-4) or GFP alone as a control were generated (Fig. 5M, lane 6) (Tillotson et al., 2017). The expression of recombinant mouse Mecp2-GFP or GFP was confirmed by immunoprecipitation with GFP antibodies followed by GFP immunoblotting (Fig. 5M compare lanes 2 with 3-4). Expression of Mecp2-GFP in *MECP2*-null cells increased respiration as determined by minimally overlapping 95% confidence intervals (Fig. 5N). Both basal and maximal respiration were increased by expressing Mecp2-GFP in two independent clones as compared to *MECP2*-null cells expressing GFP alone (Fig. 5O-P). The results indicate that mitochondrial respiration is modulated by the expression of Mecp2 in a cell-type dependent manner.

### Consumption of Exogenously Added Pyruvate is Impaired in *MECP2* mutants

Differentiated LUHMES and HAP1 *MECP2* mutant cells have decreased mitochondrial respiration that can be increased by the expression of the *Mecp2*/*MECP2* gene. However, these cells do not change their mitochondrial carrier Mpc1 levels (Fig. 4F and H). This observation can be interpreted in two ways: first, defective respiration could be related to Mpc1 function rather than absolute Mpc1 levels in these cells. Alternatively, mutant cells could have compromised mitochondrial pyruvate utilization by mechanisms downstream of Mpc1. To discriminate between these two mechanisms, the ability of *Mecp2*/*MECP2* mutant cells to use exogenously added pyruvate as the only mitochondrial fuel source was assessed. As a benchmark, *Mpc1* mutant cells preferentially compromise their FCCP-induced maximal respiration when pyruvate is the only fuel source exogenously offered to cells. This defect can be bypassed by offering glutamine as a mitochondrial fuel to Mpc1 mutant cells (Fig. 6A) (Bricker et al., 2012; Yang et al., 2014; Zhang et al., 2020).

**Figure 6.**
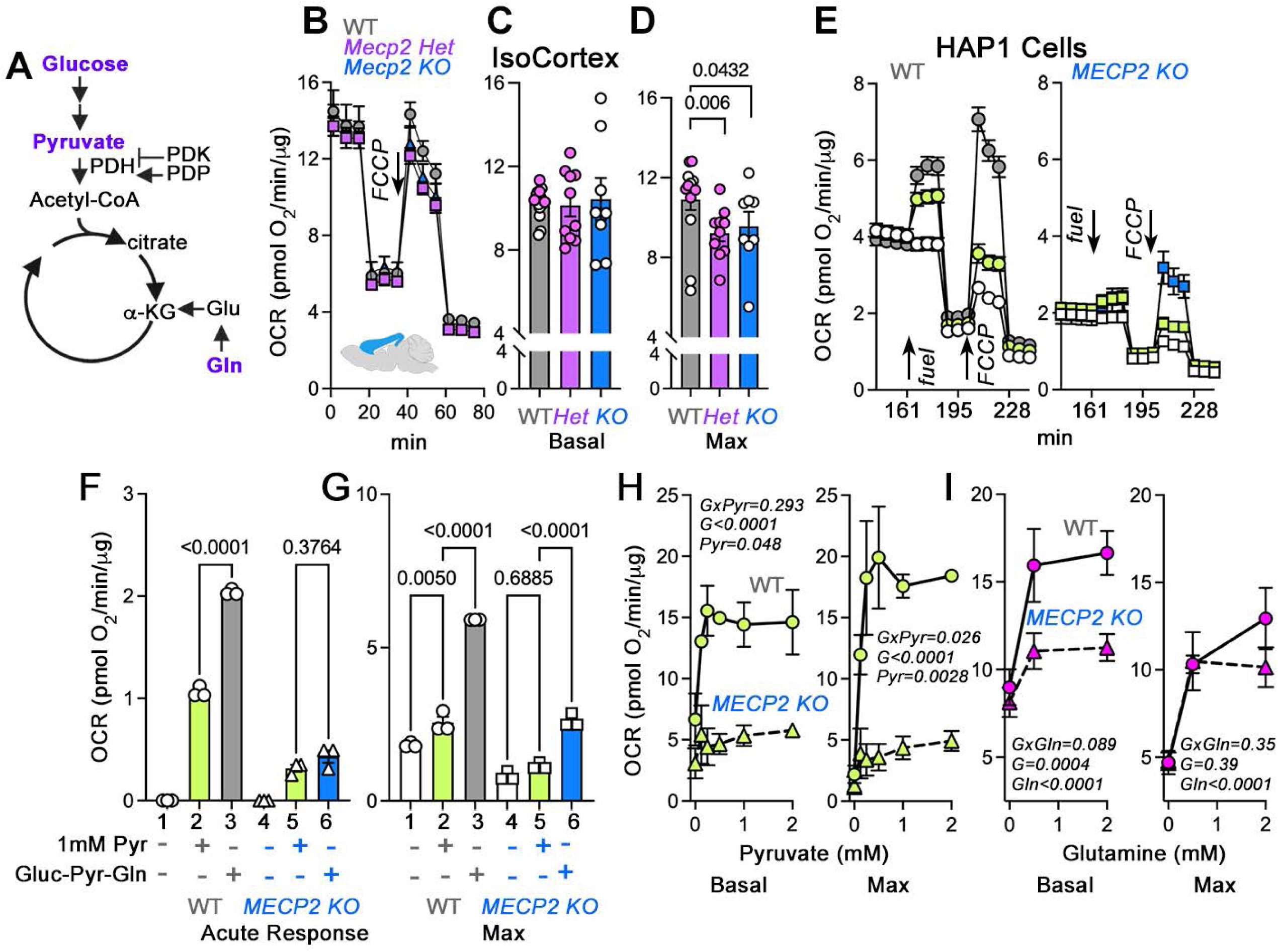
Exogenous Pyruvate and Glutamine Differentially Support Mitochondrial Respiration in *Mecp2^tm1.1Bird/y^* Isocortex Primary Cultures and *MECP2* Mutant Cells. **A.** Diagram of key glycolysis and Krebs cycle steps analyzed. Purple font indicates metabolites experimentally manipulated**. B-D.** Primary cultured isocortex neurons from postnatal 1 cultured for 14 days were subjected to a Seahorse stress test in the presence media with 1 mM pyruvate as the only fuel source. **C.** Depicts basal respiration and **D** presents maximal respiration elicited after FCCP addition (arrow). Each symbol corresponds to an independent animal, pink symbols mark female animals. p values were obtained with two-sided permutation t-test with 5000 bootstrap samples**. E** to **G** show mitochondrial stress test, acute response to fuel addition, and maximal respiration in wild type and *MECP2*-null HAP1 cells in media without fuel (white symbols) complete media (gray and blue symbols), or media with 1 mM pyruvate (green symbols). For E, inflection after data point 4 marks addition of the fuel source to basal media. One-way ANOVA followed by Bonferroni corrections. Each symbol represents an independent biological replicate. Arrows indicate fuel and FCCP additions. **H-I** depict mitochondrial basal, and maximal respiration in wild type and *MECP2*-null HAP1 cells fed media in the absence or presence of increasing concentrations of either pyruvate (H) or glutamine (I). For H n=2 and for I n=8. Two-way ANOVA where factors are genotype (*G*) and fuel, followed by Benjamini, Krieger and Yekutieli corrections. B and E, inflections after datapoint 3 or 7 respectively, mark addition of oligomycin and drop after datapoint 9 or 13 marks addition of rotenone and antimycin. All data show average ± SEM.

Mitochondrial respiration was first measured in wild type, *Mecp2^tm1.1Bird/y^* and mosaic female *Mecp2^tm1.1Bird/+^* isocortical cultures by Seahorse respirometry in media with pyruvate as the only source of fuel. A small but significant decrease in the maximal respiratory capacity in both pyruvate-fed primary neuronal cultures of *Mecp2*-null male and *Mecp2* mosaic female mice was documented (Fig. 6B-D). The small dynamic range of this pyruvate-fed isocortex phenotype prevented further study of fuel preferences by these cells. We therefore assessed HAP1 cells (Fig. 6E-I) as their MECP2 mutant proteome has the biggest overlap with the *Mecp2* mutant mouse brain proteome (Fig. S7B). Respiration in intact cells incubated in glucose-, pyruvate- and glutamine-free (fuel-free) media was assessed. Pyruvate or a mix of glucose, pyruvate and glutamine (complete media) were then acutely added as fuel sources (Figs 6E-I). Compared to wild type cells, *MECP2* mutant cells showed impaired respiration irrespective of the fuel source (Fig. 6E). Addition of pyruvate increased respiration of wild type cells as compared to fuel-free media (Fig. 6F compare columns 1-2). However, this response was half the magnitude observed in wild type cells fed complete media (Fig. 6F compare columns 2-3). In contrast, there was no difference in the magnitude of the increase in respiration rate when *MECP2*-null cells were acutely fed pyruvate-containing or complete media (Fig. 6F compare columns 5-6). Addition of FCCP increased respiration in wild type cells fed pyruvate-only or complete-media (Fig. 6G compare columns 1 with 2-3). However, *MECP2* mutant cells failed to undergo an FCCP-dependent increase in mitochondrial respiration when the only source of fuel was pyruvate (Fig. 6G compare columns 4-5). These mutant cells increased their FCCP-dependent respiration when fed complete media (Fig. 6G compare columns 4-5 with 6). This suggest that fuels other than pyruvate, like glutamine, mediate the increased FCCP-induced maximal respiration observed in mutant cells (Fig. 6A).

We compared basal and maximal respiration in wild type and *MECP2* mutant cells where the only source of fuel was either increasing concentrations of pyruvate or glutamine (Fig. 6H-I). Basal and maximal respiration increased with increasing concentrations of pyruvate in wild type cells. However, *MECP2* mutants failed to increase either respiratory parameter when fed increasing concentrations of pyruvate (Fig. 6H compare circles and triangles). In contrast to cells fed pyruvate, maximal respiration was similar between wild type and mutant cells when the only source of extracellular fuel was glutamine (Fig. 6I compare circles and triangles). These results demonstrate that HAP1 *MECP2* mutant cells have a defect in the utilization of exogenously added pyruvate when mitochondria are uncoupled with FCCP. These findings support the conclusion that mitochondria from *MECP2* mutant cells have a defect utilizing pyruvate that phenocopies Mpc1 gene defects.

### Mitochondrial Consumption of Pyruvate is Impaired in *MECP2* mutants

We next assessed whether *MECP2* mutant cells consume glycolysis-generated endogenous pyruvate by measuring respiration after a galactose switch, a condition where glucose is replaced by galactose in media. This switch halts glycolysis, thus the endogenous production of pyruvate from glucose, and renders cell ATP supply dependent solely on mitochondrial respiration (Fig. 7A) (Arroyo et al., 2016; Balsa et al., 2019; Marroquin et al., 2007). Under a galactose switch, cells use glutamine as a major fuel source for respiration (Fig. 7A) (Balsa et al., 2019). We reasoned that if *MECP2* mutant cells have impaired mitochondrial pyruvate utilization but normal glutamine utilization (Fig. 6H-I), then *MECP2* respiration should be similar to wild type respiration only in galactose-containing media. Seahorse respirometry was performed in wild type and mutant cells in media containing either glucose, pyruvate, and glutamine (complete glucose media) or the same formulation where glucose was replaced by galactose (complete galactose media) as well as controls for both media formulations that excluded glutamine (Fig. 7B-E). Complete galactose media changed the stress test respiration profile in both cell types in a glutamine-sensitive manner (compare dark and light symbols, Fig. 7B-C). The decreased respiration phenotype observed in *MECP2*-null cells was overt in complete glucose media, but wild type and mutant respiration became indistinguishable in complete galactose media (Fig. 7B-C).

**Figure 7.**
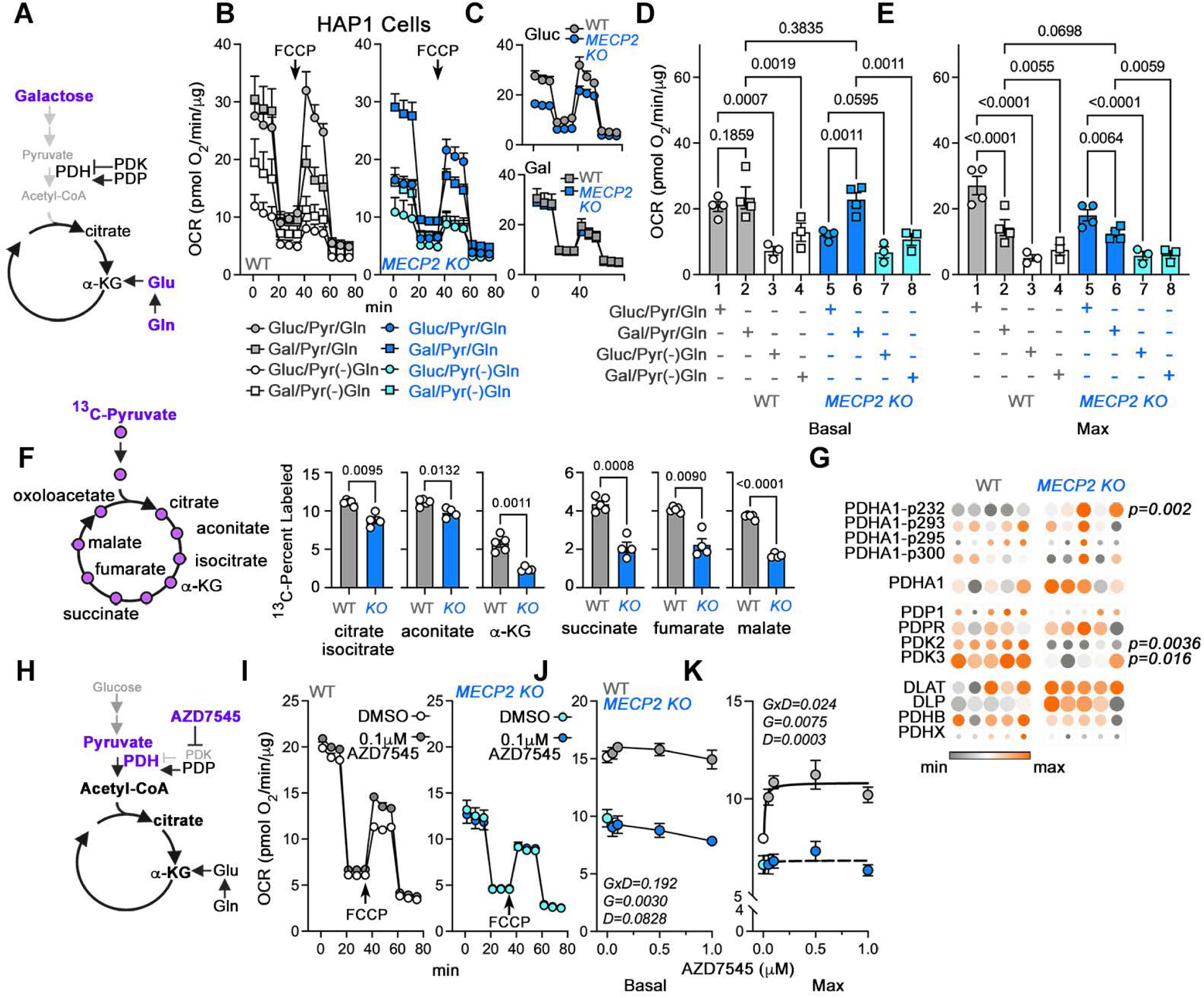
The Utilization of Endogenous Pyruvate for Mitochondrial Respiration is Selectively Impaired in *MECP2* Mutant Cells. **A.** Diagram of key glycolysis and Krebs cycle steps analyzed. Purple font indicates metabolites experimentally manipulated**. B-E.** show mitochondrial stress test, basal, and maximal respiration in wild type (white and gray symbols) and *MECP2*-null HAP1 cells (blue and teal symbols) in glucose and galactose complete media with or without glutamine [(−)Gln]. **C**. Compares complete glucose and galactose media in wild type and mutant cells shown in B. **D-E**, each symbol represents an independent biological replicate. One-way ANOVA followed by Benjamini, Krieger and Yekutieli corrections. **F,** conversion of ^13^C_3_-pyruvate into metabolites of the Krebs cycle in wild type and *MECP2*-null HAP1 cells (blue bars), n=5. **G,** heat map of the proteome and phosphoproteome log_2_ normalized expression levels for subunits of the pyruvate dehydrogenase complex, pyruvate dehydrogenase kinases and subunits of the pyruvate dehydrogenase phosphatase in wild type and *MECP2*-null HAP1 cells. Each column is an independent biological replicate. For **F** and **G**, p values were obtained with two-sided permutation t-test with 5000 bootstrap samples. **H**. Diagram of key glycolysis and Krebs cycle steps analyzed. Purple font indicates metabolites experimentally manipulated or drugs added and their target. **I**, mitochondrial stress test, **J**, basal, and **K**, maximal respiration in wild type and *MECP2*-null HAP1 cells fed 1mM pyruvate media in the presence of vehicle or increasing concentrations of AZD7545. **I** to **K**, n=4. **I** and **J**, mixed-effect analysis where factors are genotype (*G*) and drug (*D*), followed by Šídák’s multiple comparisons test. B, C and I inflections after datapoint 3 marks addition of oligomycin and drop after datapoint 9 marks addition of rotenone and antimycin. All data show average ± SEM.

Wild type basal respiration was comparable in complete glucose and galactose media (Fig. 7D compare columns 1-2). In contrast, *MECP2* mutant cells increased their basal respiration to wild type levels in complete galactose media as compared to complete glucose media (Fig. 7D compare columns 2 and 6). Basal respiration was decreased to similar levels in glutamine-free media for both genotypes (Fig. 7D compare columns 1-2 with 3-4, 5-6 with 7-8). The effect of the galactose switch and its glutamine-dependency were also evident when measuring FCCP-induced maximal respiration (Fig. 7E). Wild type maximal respiration decreased by 50% in complete galactose media as compared to glucose complete media, reflecting a requirement of endogenous pyruvate (Fig. 7E compare columns 1-2). In contrast, the FCCP-induced maximal respiration drop by only 30% in complete galactose media but to the same level observed in wild type galactose media-treated cells (Fig. 7E compare columns 2 and 6). These respiratory parameters were similarly dependent on the presence of glutamine in the media in both genotypes (Fig. 7E compare columns 1-2 with 3-4, 5-6 with 7-8). We directly measured the utilization of pyruvate by mitochondrial by measuring the conversion of ^13^C_3_-pyruvate to ^13^C-labeled Krebs cycle intermediaries (Fig. 7F). Mutant MECP2 cells showed significantly decreased conversion of labeled pyruvate into Krebs cycle carboxylic acids as compared to wild type (Fig. 7F). We conclude that *MECP2* deficient mitochondria have impaired utilization of endogenous pyruvate as a fuel source while maintaining normal consumption of glutamine for respiration.

We assessed alternative mechanisms that could account for the impaired pyruvate utilization in *MECP2* mutant cells, other than the levels of the pyruvate transporter Mcp1. The pyruvate dehydrogenase complex (PDH, Fig. 7A, G and H) is a key regulatory hub in carbohydrate metabolism that links pyruvate produced by glycolysis to the mitochondrial tricarboxylic acid cycle via its product acetyl-CoA, which is turn into citrate (Fig. 7A and H) (Gray et al., 2014). The activity of PDH is inhibited by phosphorylation of its PDHA1 subunit, a process under dynamic control of mitochondrial kinases (PDK, Fig. 7 A, G and H) and phosphatases (PDP, Fig. 7 A, G and H) (Patel et al., 2014). The abundance of PDH complex subunits was quantified by mass spectrometry (CORUM complex ID 1375; PDH1, DLAT, DLP, PDHB, and PDHX) (Hiromasa et al., 2004), PDH1 phosphopeptides, PDH kinases (PDK2 and 3), and the PDH phosphatase catalytic and regulatory subunits (PDP1 and PDPR, Fig. 7G) (Gray et al., 2014). A significant increase in the phosphorylation of PDHA1 at position 232 was identified without changes in the total abundance of PDHA1 or other subunits of the PDH complex (Fig. 7G). PDHA1 phosphorylation at position 232 inhibits PDH enzymatic activity (Korotchkina and Patel, 2001; Rardin et al., 2009). The increased PDHA1 phosphorylation could not be explained either by elevated steady-state levels of kinases or by decreased phosphatase levels (Fig. 7G). These findings suggest that changes in the enzymatic activity of either the kinase or the phosphatase in *MECP2*-null cells as a potential mechanism for PDHA1 phosphorylation at position 232. To discriminate between an increased kinase activity or a decreased phosphatase activity regulating PDHA1 phosphorylation effect of the PDK inhibitor AZD7545 (Kato et al., 2007) was assessed by Seahorse respirometry with pyruvate as the only fuel source (Fig. 7H-K). AZD7545 increased FCCP-induced maximal respiration in wild type cells, suggesting that a steady-state kinase activity acts as a brake in the utilization of pyruvate (Fig. 7I and K). In contrast, AZD7545 did not modify the FCCP-induced maximal respiration in *MECP2* mutant, a finding consistent with a model where an already decreased phosphatase activity hinders the utilization of pyruvate by the mitochondria in *MECP2*-deficient cells (Fig. 7J and K). We conclude that the *MECP2* gene defect impairs pyruvate mitochondrial utilization through a post-translational mechanism that inhibits the PDH complex.

## Discussion

To date, Rett syndrome research has primarily focused on neuronal mechanisms of disease with an emphasis on transcriptome studies. However, MECP2 is expressed in most organs with protein levels similar to those in the brain (Schmidt et al., 2018). This finding suggests that mutations in this gene have systemic effects that contribute to disease burden (Borloz et al., 2021; Kyle et al., 2018; Vashi and Justice, 2019). We comprehensively assessed steady-state levels of transcripts and proteins in multiple organs and brain regions from *Mecp2*-null mice to determine the extent to which mutations in this gene impact the whole organism. Our findings reveal that the transcriptomes and proteomes of cortex, liver, kidney and skeletal muscle undergo extensive modifications in *Mecp2^tm1.1Bird/y^* animals that occur before the appearance of overt phenotypes and the onset of mortality. Transcriptome and proteome modifications are most pronounced in brain cortex, liver, kidney and skeletal muscle as compared to hippocampus and striatum, whose transcriptomes and proteomes are modestly affected in *Mecp2* mutants. *Mecp2*-sensitive transcriptomes, but more prominently proteomes, are enriched in metabolic annotated gene products across tissues (Kyle et al., 2018). The metabolic annotated genes preponderantly represent lipid metabolism and mitochondrial pathways. Importantly, the proteome of the most affected brain region, the *Mecp2* mutant cortex, is enriched for both synaptic and mitochondrial annotated proteins by two-fold, suggesting the potential for comparable contribution of synaptic and mitochondrial processes to the genesis of Rett syndrome brain phenotypes. These findings are in agreement with recent findings in iPSC-derived human MECP2 mutant astrocytes (Sun et al., 2023; Tomasello et al., 2023). We conclude that mutations in *Mecp2* cause a systemic disorder affecting lipid and mitochondrial metabolism.

Our lipid and mitochondrial metabolic findings suggest a dysregulation of pyruvate-acetyl-CoA metabolism in *Mecp2*/*MECP2* tissues and cells. Acetyl-CoA synthesis is tied to the entry and metabolization of pyruvate by mitochondria, which we show is perturbed in *MECP2* mutant cells (Guertin and Wellen, 2023). Among the mutant tissues whose proteomes revealed altered metabolic pathways, the *Mecp2*-mutant liver proteome stands-out. We note that 10 of the 17 enzymes in the cholesterol synthesis pathway have decreased steady-state protein levels in *Mecp2* mutant liver. The robustness of this finding is supported by replication in an independent cohort of animals (data not shown). The cholesterol synthesis pathway begins with acetyl-CoA as a primary substrate. Among these 10 cholesterol synthesis enzymes down-regulated in the *Mecp2* liver, we note that squalene epoxidase (*Sqle*) encodes a rate-limiting enzymes in the cholesterol synthesis pathway (Miller and Auchus, 2011). Importantly, single copy mutation of the *Sqle* gene rescues *Mecp2*-null phenotypes revealing the centrality of this metabolic pathway to a *Mecp2*-null mouse model and the capacity of our proteome studies to capture this important disease biology (Buchovecky et al., 2013; Enikanolaiye et al., 2020). Other lipid metabolism proteins that generate acetyl-CoA are increased in *Mecp2* mutant liver. Acss3, a mitochondrial propionate-CoA ligase, catalyzes the synthesis of acetyl-CoA from short fatty acids (Yoshimura et al., 2017). Increased expression of Acss3 may be adaptive and normalize availability of acetyl-CoA. This concept is supported by a genetic suppressor screen demonstrating that mutations in acetoacetate-CoA ligase (*Aacs*) suppress phenotypes in *Mecp2^tm1.1Bird/y^* mice (Enikanolaiye et al., 2020). *Aacs* encodes a acetoacetyl-CoA synthetase that takes acetyl-CoA to funnel it to the cholesterol synthesis pathway (Ohgami et al., 2003). The hypothesis that a pyruvate-acetyl-CoA axis is regulated by the Mecp2 protein is supported by the results in cerebrospinal fluid from *Mecp2*-null mouse documenting changes in the levels of mitochondrial factors that metabolize pyruvate (Zlatic et al., 2022). *Mecp2* deficiency has also been reported to decrease levels of pyruvate dehydrogenase E1 subunit beta, and levels of this subunit are restored when *Mecp2* is re-expressed (Cortelazzo et al., 2020). Additional evidence for a dysregulated pyruvate-acetyl-CoA axis is the observation that pyruvate increases in cortex of *Mecp2^tm1.1Bird/y^* mice (Golubiani et al., 2021). Elevated pyruvate levels have also been documented in the CSF of Rett patients (Budden et al., 1990; Matsuishi et al., 1992; Matsuishi et al., 1994) and in plasma (Neul et al., 2020). Similarly, human astrocytes carrying mutations in *MECP2* have decreased levels of intracellular citrate (Sun et al., 2023; Tomasello et al., 2023). These consistent findings support the idea that impaired utilization of pyruvate is a common Rett syndrome phenotype.

Our findings in *Mecp2* mutant liver cholesterol pathways differ from those previously reported. However, there are differences in study designs that may explain this apparent discrepancy (Buchovecky et al., 2013; Enikanolaiye et al., 2020; Kyle et al., 2016). First, we analyzed animals at 45 days rather than 56 days. This age difference is important as *Mecp2^tm1.1Bird/y^* mice aged to 56 days experience a mortality rate near 50% and most mutants experience neurological symptoms (Guy et al., 2001). Second, we studied *Mecp2* mutant livers that were devoid of overt fatty liver pathology. This suggest that liver pathology in Rett syndrome models evolves with age and may be initiated by a mechanism that indirectly involves transcriptional control by Mecp2 before the onset of fatty liver (Kyle et al., 2016).

Our results expand the evidence indicating that Rett syndrome is a systemic metabolic disease that affects neurodevelopment (Kyle et al., 2018). Considering the Rett syndrome, and other disorders of neurodevelopment, as systemic diseases provides the conceptual basis to expand biomarker and therapeutic discovery (Faundez et al., 2019). Specifically, other more accessible tissues have potential to serve as a proxy for brain disease mechanisms and therapies. For example, *Mecp2* defective cholesterol synthesis pathways in liver have a parallel in brain where de novo synthesis of cholesterol is diminished (Buchovecky et al., 2013). Brain and liver defects in cholesterol synthesis may be linked to changes in the content of HDL lipoproteins in the cerebrospinal fluid of *Mecp2* mutant animal and neuronal models as well as in *Mecp2^tm1.1Bird/y^* mouse plasma (unpublished results) (Zlatic et al., 2022). These findings suggest that a plasma biomarker assessing a Rett syndrome liver phenotype has potential to correlate with similar processes in brain. Similarly, therapeutics that ameliorate peripheral tissue phenotypes could act as a proxy for the effectiveness of those therapies in brain. Take for example the therapeutic use of short fatty acid-rich or ketogenic diets that bypass the need of pyruvate as a mitochondrial fuel in Rett syndrome. These nutrients can improve *Mecp2* mouse mutant behaviors, survival, and modify liver and muscle metabolic phenotypes (Park et al., 2014). Similarly, ketogenic dietary interventions have been reported to improve neurological and behavioral phenotypes in Rett patient case reports. The effectiveness of this treatment may be more widespread and awaits results from ongoing [clinical trials](https://clinicaltrials.gov/ct2/show/NCT02696044) (Haas et al., 1986; Liebhaber et al., 2003).

There are two findings that support novel mechanisms of disease. First is the variable correlation of *Mecp2*-null transcriptomes and proteomes within a tissue. The second is a selective metabolic defect in the utilization of pyruvate by *MECP2*-mutant cells yet normal utilization of glutamine as a carbon source for respiration. *Mecp2*-sensitive mRNAs and protein steady-state levels strongly correlate in liver. In contrast, changes in the transcriptome of the *Mecp2* mutant cortex do not correlate with the changes in the proteome. The simplest explanation for this lack of correlation in cortex would be technical problems with either the proteome or transcriptome assays or data processing pipelines. However, this is unlikely for several reasons. First, we performed all tissue analyses simultaneously, therefore technical problems with brain tissue should have also affected analyses performed in the other tissues. Second, we validated our transcriptome against multiple replication benchmarks. Third, we asked how well the wild type cortex transcriptome and proteome correlate to find that the Spearman correlation is 0.38. This value is within the range of similar correlations across tissues, including nervous tissue, and diverse cells (Spearman 0.24–0.65) (de Sousa Abreu et al., 2009; Jiang et al., 2020; Mohammed et al., 2021; Sharma et al., 2015; Wang et al., 2019). Finally, our proteomics shows strong ontological/pathway convergence with a previous *Mecp2* mutant mouse brain cortex proteome (Pacheco et al., 2017). Confidence in the validity of our analyses is further increased by the robustness of the replication despite considerable differences in methodology including a different *Mecp2* animal model, timing and method of tissue collection, the mass spectrometry strategy, and data processing between our study and those employed for the whole isocortex proteome (Pacheco et al., 2017). Furthermore, the ontological/pathway similarity of the mouse brain *Mecp2* cortex proteome extends to the mouse and rat cerebrospinal *Mecp2* mutant proteomes demonstrating robustness in pathway identification in *Mecp2* mutant brain (Zlatic et al., 2022). Our data suggest that the poor correlation between the *Mecp2*-sensitive brain transcriptomes and proteomes may be disease phenotype. Divergency between the *Mecp2*-sensitive brain transcript and protein levels could be explained by posttranscriptional or posttranslational mechanisms. A factor contributing to transcriptome-proteome divergence is a spatial segregation of the transcriptome from the proteome that it controls, such as dissimilar localization of the transcript and the protein it encodes at an anatomical/subcellular level (Buccitelli and Selbach, 2020; Liu et al., 2016). Transcriptomes could also be more susceptible to cell diversity than the proteome. Brain and liver have marked differences in cell type diversity but cellular type diversity is similar between skeletal muscle and liver (Tabula Muris et al., 2018). Nonetheless, muscle and liver have pronounced differences in the correlation of their transcriptome-proteome correlations. Finally, we explored whether small regulatory RNAs under the control of *Mecp2* could account for modifying transcriptome output (Cheng et al., 2014; Szulwach et al., 2010; Urdinguio et al., 2010; Wu et al., 2010). We measured the levels of microRNAs in microdissected cortex and liver were quantified as these two tissues had the most pronounced differences in their transcriptome-proteome correlations. The predicted targets of the microRNAs that have altered expression with *Mecp2* mutations only minimally overlap with cortex and liver transcript and proteins affected by the *Mecp2* mutation. These findings do not support the hypothesis that MECP2 protein indirectly regulates translation through regulation of microRNAs. It is likely that other tissue-specific regulatory factors or combinations of thereof explain *Mecp2*-mutant transcriptome-proteome divergency. Potential mechanisms include differences in RNA stability, a proteome with a delayed response to modifications of the transcriptome, as well as changes in the rate of protein synthesis and/or degradation (Buccitelli and Selbach, 2020; Liu et al., 2016). Some of the mechanisms may be intertwined, as recently demonstrated in *MECP2*-mutant iPSC-derived neurons where a global decrease of the active translatome and a decrease in mTOR-dependent protein synthesis impact proteins levels, including components of the protein degradation machinery (Rodrigues et al., 2020). Since the translatome varies across development and tissues, and is modified by the metabolic status, it reasonable to speculate that the differences observed across *Mepc2*/*MECP2* mutant tissue transcriptome-proteome correlations are determined, in part, by cell-autonomous translational, post-translational, and metabolic landscapes (Harnett et al., 2022; Ho et al., 2020; Rodrigues et al., 2020; Wang et al., 2021; Wang et al., 2020; Wu et al., 2010).

Another novel finding is the *MECP2*-mutant defect in pyruvate utilization yet normal consumption of glutamine for respiration. We think this observation reveals an impaired metabolic flexibility phenotype, a hallmark of metabolic diseases. Mitochondria modify the quality and quantity of the carbon substrate oxidized by mitochondria, a process known as metabolic flexibility (Goodpaster and Sparks, 2017; Smith et al., 2018). The plasticity in the quality and/or magnitude of mitochondrial substrates utilized is determined by the tissue functional demand, organism metabolic status (lean or obese), developmental stage, the availability of nutrients, or a pathological condition such as diabetes (Goodpaster and Sparks, 2017; Smith et al., 2018). The fetal brain relies on aerobic glycolysis but at birth it progressively transitions to consumption of oxygen and glucose reaching a maximum around postnatal age five years (A et al., 2021; Bulow et al., 2022; Goyal et al., 2014; Kuzawa et al., 2014). Pyruvate derived from glycolysis is a key metabolite whose entry to mitochondria, modulates carbon utilization preferences between glucose, aminoacids, ketones, and lipids (Jeon et al., 2021; Olson et al., 2016). In the case of neurons, pyruvate-dependent metabolic flexibility couples the activity of the synapse to the production of ATP by synaptic mitochondria (Ashrafi et al., 2020; Diaz-Garcia et al., 2021). The changes in expression of either the pyruvate carrier Mpc1 or the NAD^+^ carrier Slc25a51 in *Mecp2* mutant brains support the idea of defect defective metabolic flexibility in neural tissue. Thus, impaired metabolic flexibility could contribute to synaptic phenotypes in Rett syndrome and other neurodevelopmental disorders (Chao et al., 2007).

## Material and Methods

### Animals

All animal procedures and husbandry were approved by Emory University’s Institutional Animal Care and Use Committee. The Mecp2-null mouse model, B6.129P2(C)-Mecp2^tm1.1Bird^/J (Cat: 003890 RRID: IMSR_JAX:003890), was purchased from The Jackson Laboratory at 5-6 weeks of age and housed within Emory University’s Division of Animal Resources for about 4 days before tissue collection.

### Mouse Tissue Dissection

Mice were euthanized by CO_2_ asphyxiation and tissues dissected at 6 weeks of age. Tissue collection was done in the morning and animals were sacrificed by alternating wild type and mutant genotypes to minimize circadian rhythm impact. Liver, kidney, whole brain, and skeletal muscle from the hind quarters (gluteus maximus, rectus femoris or vastus lateralis) of the mouse was removed and washed in ice cold phosphate buffered saline. For micro-punch dissections, all tools and surfaces for dissection were prechilled. Whole brain was transferred to an aluminum coronal slicing matrix with 1.0 mm slice intervals (Zivic Cat: BSMAS001-1 and BSMYS001-1) and sliced with single edge razors. Slices were laid out on an aluminum block and a micro-punch tool kit (Stoelting Cat: 57401) with 1.00 mm punch (Stoelting Cat: 57397) was used to micro-dissect from hippocampal, audio-visual region of cortex, and striatal region. Punches from the left and right hemispheres were pooled for approximately six punches per region. The remaining cortex from all the coronal slices was collected for bulk cortical samples. All samples were flash frozen with liquid nitrogen and stored at −80°C (Barr et al., 2004).

### Mouse Primary Culture

All tools were sterilized and dissection reagents prechilled. Postnatal day 0-2 pups were euthanized by decapitation. Whole brain was removed and washed in HBSS/10mM HEPES. Cortex was dissected from the whole brain, cut into 5-6 smaller pieces and digested in prewarmed trypsin solution for 10 minutes. Cortical pieces were then washed briefly in HBSS/10 mM HEPES twice and transferred to MEM solution consisting of MEM (Sigma M4655) with 10 mM HEPES (Gibco 15630-080), 33 mM D-glucose (Sigma G8769), and 10% FBS (VWR 97068-085) solution. Tissue was triturated 8 times through a fire polished glass pipette. Suspended cells were collected and gently centrifuged for 5min at 130 RCF, 4°C. Supernatant was removed and cells were resuspended in fresh MEM/HEPES/D-glucose/FBS solution. Cells were plated onto 0.01% poly-L-lysine (Sigma P4707) and 2 µg/cm^2^ laminin (Sigma L2020) coated dishes and cultured in a humidified 37°C, 5% CO^2^ incubator. After cells had settled, approximately 2 hours, media was changed to Neurobasal (Gibco 21103-049) supplemented with 1x B27 (Gibco 17504-044) and 2 mM L-glutamine. Half the volume of media was changed out every 2-3 days thereafter until assayed at 14 days in vitro.

### Tissue Culture

Control and two clonal MECP2 knock out (2_7 and H4) LUHMES cell lines were gifts from Dr. Adrian Bird (Shah et al., 2016). Cells were passaged in Advanced DMEM/F12 (Gibco 12634-010) supplemented with 1x N2 (Gibco 17502048), 2 mM L-glutamine (Sigma G7513), and 40 µg/ml FGF (R&D Systems, 4114-TC) at 2,000,000 – 500,000 cells/75 cm^2^ flask. Three million cells/75 cm^2^ were pre-differentiated for 48 hours in differentiation media, lifted, and seeded at approximately 1 million cells/cm^2^ onto assay plates coated with poly-L-ornithine 44 µg/mL (Sigma P3655) and fibronectin 1 µg/mL (Sigma F1141) in differentiation media. Differentiation media consists of Advanced DMEM/F12 (Gibco 12634-010) supplemented with 1x N2 (Gibco 17502048), 2 mM L-glutamine (Sigma G7513), 1 mM DbcAMP (Sigma D0627) 1µg/mL tetracycline (Sigma T7660) and 2 ng/mL GDNF (R & D Systems 212-GD). MECP2 KO SHSY5Y cells were developed with Synthego. Synthego used guide RNA sequence CCACTCTGCTGAGCCCGCAG, for genome editing of MECP2 at exon 3, in Cas9-gRNA ribonucleoprotein complexes. A knock-out pool of 93% efficiency was diluted to clonal populations. Clones were confirmed by Sanger sequencing and Western blot. Cells were cultured in DMEM (Corning 10-013-CV) with 10% FBS (VWR 97068-085) at 37°C, 5% CO^2^. For differentiation, approximately 40,000 cells/cm^2^ were seeded onto plates coated with 0.01% poly-L-lysine (Sigma P4707) per manufacturer’s protocol and switched to differentiation media consisting of Neurobasal (Gibco 21103-049) supplemented with 1x B27 (Gibco 17504-044), 500 µM L-glutamine (Sigma G7513), and 1 mM DbcAMP (Sigma D0260) for 3 days and then assayed. MECP2 knock out HAP1 cells from Horizon Discovery (Cat: HZGHC001102c010, RRID: CVCL_SX72) were cultured in IMDM ((Lonza, 12-722F) supplemented with 10% FBS (VWR 97068-085) in a humidified, 37°C, 10% CO^2^ incubator. For MECP2-GFP rescue of HAP1 cells, pEGFP-N1_MeCP2(WT) was a gift from Adrian Bird (Addgene plasmid #110186; RRID:Addgene_110186), GFP only vector was created by digesting pEGFP-N1_MeCP2(WT) with BamHI and BglII to remove the MECP2 insert and ligated with T4 DNA ligase. Plasmids were transfected to HAP1 cells using FUGene according to manufactures protocol in Optimem Media (Gibco 31985070). Transfected cells were selected with G418 selection 600 µg/mL in growth media 48 hours after transfection media was added. Transfected cells were diluted to clonal populations and expanded for assay. For Galactose switch assays, 24 hours before assay, HAP1 cells were incubated in a basal DMEM (Gibco A14430-01) supplemented with 10% dialyzed FBS (Hyclone SH30079.03), 1 mM sodium pyruvate, 2 mM L-glutamine, 96 nM sodium selenite, non-essential amino acids, and either 10 mM glucose or 10 mM galactose. In the case of L-glutamine exclusion, the L-glutamine was not added to either the glucose or galactose version of the media.

### Seahorse Oximetry

Seahorse XFe96 FluxPaks (Agilent 102416-100) were hydrated at 37°C for approximately 24 hours in Seahorse XF Calibrant solution (Agilent 100840-000). Cells were plated on Agilent Seahorse XF96 V3-PS Microplates (Agilent 101085-004) approximately 24 hours before stress tests. Seahorse stress test media consisted of Seahorse XF base medium (Agilent 103334-100) or Seahorse XF DMEM base medium pH 7.4 (Agilent 103575-100) supplemented with 1 mM sodium pyruvate (Sigma S8636), 2 mM L-glutamine (Sigma G7513), and 10 mM D-glucose (Sigma G8769), or 10 mM galactose. For AZD7545 (Selleckchem) experiments, AZD7545 was added to seahorse assay media at 10 µM, 5 µM, 2 µM, 1 µM, 0.5 µM, 0.1 µM, and 0.05 µM AZD7545. Figures and the table below describe the exact components for each experiment. On the day of analysis, cells were washed twice in 37°C warmed Seahorse analysis medium and then the volume was brought to180 µL one hour prior to assay and incubated at 37°C in the absence of CO^2^ injection. For AZD7545 treatment, warmed media was added 2 hours prior to assay start time. In the case of nutrient starvation, warmed media was added and the assay was immediately started. The mitochondrial stress test was run as per Agilent Seahorse protocols with Seahorse Wave Software (Agilent). Seahorse Wave Software data collection was as per manufactures conditions. For standard mitochondrial stress tests, oxygen consumption rates and extracellular acidification rates were collected 3 times for each phase of the stress test with a 3-minute mix step followed by 3 minutes of measurement. FluxPak ports were loaded with stress test drugs at a 10x concentration of the final well concentration. Stress tests consisted of Oligomycin A (Sigma 75351), Carbonyl cyanide 4-(trifluoromethoxy) phenylhydrazone, FCCP (Sigma C2920), and Rotenone (Sigma R8875) and Antimycin A (Sigma A8674) at injections a, b, and c respectively. For nutrient starvation stress tests, the basal measurements were collected 40 times (about 2.5 hours) with 3-minute mix steps followed by 3 measurements with 3-minute mix steps after each injection. Well concentrations of stress test drugs were cell type dependent and described below. All seahorse microplates were normalized by total protein using the Pierce BCA Protein Assay Kit (Thermo 23227) as per protocol with a BSA protein standard. The BCA assay absorbance was read by a Biotek microplate reader using Gen5 software. Seahorse oximetry data analysis was done with Agilent Wave Software Report Generator and Microsoft Excel.

Mouse primary culture: 20,000 cells/well, 2.0 µM oligomycin, 0.5 µM FCCP, 1.0 µM rotenone, 1.0 µM antimycin A, 17.5 mM D-glucose

LUHMES: 40,000 cells/well, 2.0 µM oligomycin, 0.25 µM FCCP, 1.0 µM rotenone, 1.0 µM antimycin A HAP1: 40,0000 cells/well, 2.0 µM oligomycin, 0.125 µM FCCP, 1.0 µM rotenone, 1.0 µM antimycin A

SHSY5Y: 30,000 cells/well, 2.0 µM oligomycin, 0.25 µM FCCP, 1.0 µM rotenone, 1.0 µM antimycin A

| Figure | Seahorse Stress Test Assay Media Components |
| --- | --- |
| 6A-L,<br>N-P<br>7B-D | Seahorse XF DMEM base medium, 1 mM sodium pyruvate 2 mM L-glutamine, 10 mM D-glucose |
| 7E-G | Seahorse XF base medium |
|  | Seahorse XF base medium, 1 mM sodium pyruvate |
|  | Seahorse XF base medium, 1 mM sodium pyruvate, 2 mM L-glutamine, 10 mM D-glucose |
| 7H | Seahorse XF DMEM base medium, and 0 mM, 0.125 mM, 0.25 mM, 0.5 mM, 1.0 mM, or 2.0 mM sodium pyruvate |
| 7 I | Seahorse XF DMEM base medium, and 0 mM, 0.5 mM or 2 mM L-glutamine |
| 8B-E | Seahorse XF DMEM base medium, 1 mM sodium pyruvate, 2 mM L-glutamine, 10 mM D-glucose |
|  | Seahorse XF DMEM base medium, 1 mM sodium pyruvate, 2 mM L-glutamine, 10 mM galactose |
|  | Seahorse XF DMEM base medium, 1 mM sodium pyruvate, 10 mM D-glucose |
|  | Seahorse XF DMEM base medium, 1mM sodium pyruvate, 10 mM D-galactose |
| 8H-J | Seahorse XF base medium, 1 mM sodium pyruvate, 2 mM L-glutamine, 10 mM D-glucose |

### Resipher Oximetry

HAP1 Control and MECP2 Knock-out cells were seeded at 2,000 cells per well in a NUNC 96-well microplate (Thermo Scientific, Cat# 269787) with 200 µL standard growth media (above) and cultured overnight at 5% CO_2_, 37°C. SHSY-5Y Control and MECP2 KO cells were seeded at 40,000 cells per well on poly-l-lysine coated plates and incubated in 200 µL differentiation media (above) for 3 days at 5% CO_2_, 37°C. On the day of the experiment, the Lucid O_2_ platinum sensing probe array and Resipher device (Lucid Scientific, Inc) were pre-warmed in the 37°C incubator. In the 96-well cell plate, 75% of culture media was replaced with fresh pre-warmed growth media (HAP1) or differentiation media (SHSY5Y), the sensing probes were placed in the wells, and Resipher device was attached to the sensing probes. Resipher Oxygen Consumption calculations were gathered for over at least 48 hours in a 5% CO_2_, 37°C environment at which point rotenone and antimycin A were added to the wells at a concentration of 1µM. In order to account for cell replication, cell plates were made in triplicate and cell counts or protein concentrations were taken from these replicate plates when the Resipher experiment started, approximately 24 hours into the experiment, when rotenone/antimycin were added and/or from the actual Resipher experiment plate at the end of the experiment. Cell counts were made in HAP1 cell plates by removing culture media, adding 10mM EDTA in PBS to release cells from the plate, incubating on a rocker at 4°C for 10 minutes, and taking counts from cells suspended in solution. Protein concentration was determined in SHSY5Y cell wells by removing culture media, washing three times with PBS containing 0.1 mM CaCl_2_ and 1.0 mM MgCl_2_, and using the Pierce BCA Assay Kit (Thermo Scientific, Cat# 23227) according to manufactures specifications. Absorbance was read on a Biotek plate reader with Biotek Gen5 software. Estimated cell numbers were calculated deriving the doubling time using the formula N_t_=N_0_2exp(t/T_d_) where N_0_ is the initial cell number, N_t_ is the cell number at time t and T_d_ is the doubling time. To assess the effect of genotype on respiration and cell number we integrated the area under the curve using Prism Version 10.0.1 (170).

### Cell Lysis, SDS-PAGE, Western Blotting, and Silver Stain

For cellular lysis, cells were cultured as above, washed twice in phosphate buffered saline (Corning 21-040-CV) containing 0.1 mM CaCl_2_ and 1.0 mM MgCl_2_ and lysed in 8 urea with Complete protease inhibitor (Roche 11245200) and/or Halt phosphatase inhibitor (Pierce A32957) and incubated on ice for 30 minutes. Lysis was sonicated for 10 short bursts at 20% amplitude to shear DNA. Lysis was then centrifuged at 20,000 g for 10 minutes at 4°C to clarify any insolubilized tissue. Total protein content was determined by Bradford (BioRad 5000006) or BCA (Pierce 23227) assays as per manufactures’ protocols. For SDS-PAGE, lysis was reduced and denatured in Laemmli buffer containing SDS and 2-mercaptoethanol and heated for 5 minutes at 75°C. Equal micrograms of samples were loaded onto 4% - 20% Criterion gels (BioRad 5671094) for electrophoresis and used for silver staining or transferred to PVDF (Millipore IPFL00010) using the semi-dry transfer method for western blotting. For silver stain, gels were fixed overnight in 50% methanol, 12% acetic acid, and 0.02% formalin. The following day the gel was washed three times, 20 minutes each, with 50% ethanol followed by 1-minute soak in 2 mM sodium thiosulfate. Gels were then washed again with three quick washes in water and soaked in a 12 mM silver nitrate, 0.03% formalin solution for 20 minutes. After another 2 quick washes, gel was developed in 1.2% sodium carbonate, 0.004% formalin, 7.6 mM sodium thiosulfate. For western blots, membranes were blocked in Tris buffered saline containing 0.05% Triton X-100 (TBST) and 5% non-fat milk for 30 minutes at room temperature. Membranes were then rinsed of blocking solution and incubated overnight with primary antibody diluted in antibody base solution (PBS, 3% bovine serum albumin, 0.2% sodium azide). Membranes were washed in TBST. HRP conjugated secondary antibodies were diluted 1:5000 in blocking solution and incubated with membranes for 30 minutes at room temperature. Washed membranes were then exposed to Amersham Hyperfilm ECL (GE Healthcare 28906839) with Western Lightning Plus ECL reagent (Perkin Elmer NEL105001EA). Densitometry was quantified with Fiji Image J software.

Primary Antibodies: MPC1 1:250 (Sigma HPA045119), MPC2 1:500 (Protein Tech Group 20049-1-AP), SLC25A1 1:500 (Protein Tech Group 15235-1-AP), MECP2 1:500 (Invitrogen PA-1-888), Actb 1:500 (Sigma, A5451), GFP 1:1000 (Synaptic Systems 132002)

Secondary Antibodies: HRP anti Mouse 1:5,000 (Invitrogen A10668), HRP anti rabbit 1:5,000 (Invitrogen G21234)

### Immunoprecipitation

For immunoprecipitation, Dynabeads Sheep anti-Mouse IgG (Invitrogen 11031) magnetic beads (30 µL) were incubated at room temperature for 2 hours with 1 µg mouse monoclonal anti-GFP (Invitrogen 3E6 A-11120), washed thrice with Wash Buffer (150 mM NaCl, 10 mM HEPES, 1 mM EGTA, and 0.1 mM MgCl_2_, pH 7.4 with 0.1% Triton X-100) and then incubated with 350 µg protein in 1:1 8M urea with Complete protease inhibitor (Roche 11245200) sample lysis:Buffer A/0.5% Triton X-100 (150 mM NaCl, 10 mM HEPES, 1 mM EGTA, and 0.1 mM MgCl_2_, with 0.5% Triton X-100 and Complete protease inhibitor (Roche 11245200)). After sample had incubated with beads for 4 hours at 4°C, the beads were washed 6 times with Wash Buffer, and proteins eluted from beads with Laemmli buffer. Samples were then resolved by SDS-PAGE followed by western blot as described above.

### RNAseq

Trizol RNA extraction, library construction, and sequencing were performed by BGI. Quality control of total RNA quality and quantity was done on the Agilent 2100 Bio analyzer (Agilent RNA 6000 Nano Kit, Cat# 5067-1511) QC: RNA concentration, RIN value,28S/18S and the fragment length distribution. For library generation, poly-T oligo-attached magnetic beads were used to isolate poly-A containing mRNAs. Poly-A mRNA was fragmented using divalent cations under elevated temperature. RNA fragments were copied into first strand cDNA with reverse transcriptase and random primers. Second strand cDNA synthesis was done using DNA Polymerase I and RNase H. cDNA fragments underwent addition of a single ‘A’ base followed by ligation of the adapter. The products were isolated and enriched by PCR amplification. PCR yield was quantified by Qubit. Samples were pooled together to make a single strand DNA circle, giving the final library. DNA nanoballs were generated with the single strand DNA circle by rolling circle replication to increase fluorescent signals during sequencing. DNA nanoballs were loaded into the patterned nanoarrays. Pair-end reads of 100 bp were read on a BGISEQ-500 platform for data analysis. The BGISEQ-500 platform combined the DNA nanoball-based nanoarrays and stepwise sequencing using Combinational Probe-Anchor Synthesis Sequencing Method.

For sequencing analysis, FastQC was used to remove samples of poor quality(Andrews, 2012). We then used the usegalaxy.org server where we were uploaded sequencing reads for analysis (Afgan et al., 2018). The Galaxy server (v. 21.01) running Hisat2 (Galaxy Version 2.1.0+galaxy7), FeatureCounts (Galaxy Version 2.0.1), and Deseq2 (Galaxy Version 2.11.40.6+galaxy1) was used to map sequence reads (Kim et al., 2015; Liao et al., 2014; Love et al., 2014). FeatureCounts files and raw files are available at GEO with accession GSE140054 and compiled file is GSE140054_AllTissueFeatureCounts.txt.gz. In order to determine differential expression, the DESeq2 package was used (Love et al., 2014). We did not change the default settings and thus the statistics on determining differential expression are well outlined in: https://hbctraining.github.io/DGE_workshop_salmon/lessons/05_DGE_DESeq2_analysis2.html Default was used to estimate size factors (for normalization) : ” ‘ratio’ uses the standard median ratio method introduced in DESeq. The size factor is the median ratio of the sample over a ‘pseudosample’: for each gene, the geometric mean of all samples. We did not do any pre-filtering of zeros across conditions. For differential expression, we stuck with the default parametric fit type because it is less sensitive to a single sample that may have a large count relative to all the other samples with low counts. And we saw decreasing gene-wise dispersion estimates over the mean; thus, a parametric fit was used for comparing expression levels. Our adjusted p-value was calculated using with the Benjamini-Hochberg procedure controlling for the false discovery rate (FDR). Because we had more than 2 replicates per condition, DESeq2 automatically filtered genes which contain a Cook’s distance above a cutoff that is determined based on sample size and number of parameters needing to be estimated. The default cutoff determined by “the 99% quantile of the F(p,m-p) distribution (with p the number of parameters including the intercept and m number of samples). (http://bioconductor.org/packages/devel/bioc/vignettes/DESeq2/inst/doc/DESeq2.html#multi-factor-designs). DESeq2 performs independent filtering by default using the mean of normalized counts as a filter statistic. Meaning it filtered out genes unlikely to produce low p-values. Thus prior to multiple testing corrections and Wald Testing (hypothesis testing) those genes with low counts (zeros) and outliers are removed (given NA values) and then those unlikely have low p-values become NA in adj-P value column. Variance-stabilized normalized counts and regularized log-transformed counts were also generated because they were preferred for any downstream clustering or visualization. We utilized Hisat2 with the following settings: paired-end, unstranded, default settings (except for when a GTF file was used for transcript assembly). For GTF files, we used the *Mus musculus* (Mouse), Ensembl, GRCm38 build from iGenome (Illumina). The aligned SAM/BAM files were processed using Featurecounts with Default settings except we used Ensembl GRCm38 GTF file and output for DESeq2 and gene length file.

### Small RNA profiling

Total RNA was purified from the dissected tissue specimens using the Qiagen miRNeasy kit. After examining RNA quality and yield, stranded libraries were prepared with the Illumina TruSeq Small RNA Library Prep kit with the following modifications to the protocol. First, the final library amplification was done with 15 cycles PCR. Second, the PCR products were pooled into 6 sets of 6 samples each prior to loading on a 4% MetaPhor agarose gel. From these gels, a single band at approximately 150 bp was excised, which was then centrifuged at 14,000 RPM through a Gel Breaker Tube (Ist Engineering). The eluted libraries were then concentrated by ethanol precipitation according to the protocol in the Illumina manual. Libraries were sequenced on an Illumina NextSeq500 instrument with a single-end 50 bp read using 2.0 pM loading concentration with 1% PhiX spike. All Mecp2 tissue samples were run together, without any other samples in the sequencing run. Samples generated an average of 8.13 million reads/sample (SE +/− 380,033), with average Q scores of 34.66. Reads were aligned to the Mm10 mature miRbase 22 reference database using a seed length of 19 bases, allowing 1 mismatch per seed. An average of 4.30 million (+/− 313,224) alignments/sample were identified, with an average alignment Q score of 34.4. Reads were quantified and filtered to include the 80^th^ percentile of features with the highest minimum count and exclude those with a maximum count <10. The resulting 503 miRNAs were then normalized using the counts per million (CPM) method, and log2 transformed before being subjected to statistical testing and downstream analyses using Qlucore Omics Explorer Version 3.6(33) using normalized data to a variance of 1 and a mean of 0 followed by t-test comparisons without multiple corrections.

### Mouse Tissue TMT Mass Spectroscopy

Each tissue sample was homogenized in 500 μL of 8 M urea/100 mM NaHPO4, pH 8.5 with HALT protease and phosphatase inhibitor cocktail (Pierce 78440) using a Bullet Blender (Next Advance) according to manufacturer protocols. Briefly, tissue lysis was transferred to a 1.5 mL Rino tube (Next Advance) with 750 mg stainless steel beads (0.9–2 mm in diameter) and blended for 5 minute intervals, two times, at 4°C. Protein supernatants were sonicated (Sonic Dismembrator, Fisher Scientific) three times for 5 seconds, with 15 second intervals of rest, at 30% amplitude to disrupt nucleic acids, in 1.5 mL Eppendorf tubes. Protein concentration was determined by BCA method, and aliquots were frozen at −80°C. Protein homogenates (100 µg) were diluted with 50 mM NH4HCO3 to a final concentration of less than 2 M urea and treated with 1 mM dithiothreitol (DTT) at 25°C for 30 minutes, followed by 5 mM iodoacetimide (IAA) at 25°C for 30 minutes in the dark. Proteins were digested with 1:100 (w/w) lysyl endopeptidase (Wako) at 25°C for 2 hours followed by another overnight digestion with 1:50 (w/w) trypsin (Promega) at 25°C. Resulting peptides were desalted on a Sep-Pak C18 column (Waters) and dried under vacuum.

In order to compare all samples per tissue type, 10 individual samples and one composite sample were labeled using the TMT 11-plex kit (ThermoFisher 90406). Labeling was performed as previously described (Higginbotham et al., 2020; Ping et al., 2020). Briefly, each sample of 100 μg vacuum desiccated peptides was re-suspended in 100 mM TEAB buffer (100 μL). The TMT labeling reagents were equilibrated to room temperature, and anhydrous ACN (256 μL) was added to each reagent channel. Channels were gently vortexed for 5 minutes, and then 41 μL from each TMT channel was transferred to the peptide solutions and allowed to incubate for 1 hour at room temperature. Reactions were quenched with 5% (v/v) hydroxylamine (8 μL; Pierce). All 10 channels were then combined and dried by SpeedVac (LabConco) to ∼150 μL and diluted with 1 mL of 0.1% (v/v) TFA, followed by acidification to a final concentration of 1% (v/v) FA and 0.1% (v/v) TFA. Peptides were desalted with a 200 mg C18 Sep-Pak column (Waters). Each Sep-Pak column was activated with 3 mL of methanol, washed with 3 mL of 50% (v/v) ACN, and equilibrated with 2 × 3 ml of 0.1% TFA. The samples were then loaded and each column was washed with 2 × 3 mL 0.1% (v/v) TFA, followed by 2 mL of 1% (v/v) FA. Elution was performed with 2 volumes of 1.5 mL 50% (v/v) ACN. Eluates were dried to completeness.

High pH fractionation was performed essentially as described with slight modification (Ping et al., 2020). Dried samples were re-suspended in high pH loading buffer (0.07% v/v NH4OH, 0.045% v/v FA, 2% v/v ACN) and loaded onto an Agilent ZORBAX 300 Extend-C18 column (2.1 × 150 mm with 3.5-μm beads). An Agilent 1100 HPLC system was used to carry out the fractionation. Solvent A consisted of 0.0175% (v/v) NH4OH, 0.0125% (v/v) FA, and 2% (v/v) ACN; solvent B consisted of 0.0175% (v/v) NH4OH, 0.0125% (v/v) FA, and 90% (v/v) ACN. The sample elution was performed over a 58.6-minute gradient with a flow rate of 0.4 mL/minute. The gradient consisted of 100% solvent A for 2 minutes, 0–12% solvent B over 6 minutes, 12–40% over 28 minutes, 40–44% over 4 minutes, 44–60% over 5 minutes, and then held constant at 60% solvent B for 13.6 minutes. A total of 96 equal volume fractions were collected across the gradient, pooled by concatenation into 24 fractions, and dried to completeness using a vacuum centrifugation.

Each of the 24 high-pH peptide fractions was resuspended in loading buffer (0.1% FA, 0.03% TFA, 1% ACN). Peptide eluents were separated on a self-packed C18 (1.9 μm Maisch) fused silica column [25 cm × 75 μm internal diameter (ID), New Objective] by an Easy nLC 1200 (Thermo Scientific) and monitored on a Q-Exactive HFX MS (Thermo Scientific). Elution was performed over a 120-minute gradient at a rate of 300 nL/minute with buffer B ranging from 3% to 40% (buffer A: 0.1% FA in water; buffer B: 0.1% FA in 80% ACN). The mass spectrometer acquired data in positive ion mode using data-dependent acquisition with top 10 cycles. Each cycle consisted of one full MS scan followed by a maximum of 10 MS/MS. Full MS scans were collected at a resolution of 120,000 (400–1600 m/z range, 3 × 106 AGC, 100 ms maximum ion injection time). Higher energy collision-induced dissociation (HCD) MS/MS spectra were acquired at a resolution of 45,000 (1.6 m/z isolation width, 30% collision energy, 1 × 10–5 AGC target, 86-ms maximum ion time). Dynamic exclusion was set to exclude previously sequenced peaks for 20 s within a 10-ppm isolation window.

Raw TMT files were searched using Thermo’s Proteome Discoverer suite (version 2.1.1.21) with Sequest HT. The spectra were searched against a mouse Uniprot database downloaded July, 2018 (98,225 target sequences). Search parameters included 20 ppm precursor mass window, 0.05 Da product mass window, dynamic modifications methione (+15.995 Da), deamidated asparagine and glutamine (+0.984 Da), phosphorylated serine, threonine and tyrosine (+79.966 Da), and static modifications for carbamidomethyl cysteines (+57.021 Da) and N-terminal and lysine-tagged TMT (+229.26340 Da). Percolator was used filter PSMs to 0.1%. Peptides were grouped using strict parsimony and only razor and unique peptides were used for protein level quantitation. Reporter ions were quantified from MS2 scans using an integration tolerance of 20 ppm with the most confident centroid setting. Only unique and razor (i.e., parsimonious) peptides were considered for quantification. Volcano plot p values were calculated using Qlucore Omics Explorer Version 3.6(33). Data were normalized to a variance of 1 and a mean of 0 followed by t-test comparisons without multiple corrections.

### Cell Lines TMT Mass Spectrometry

Cells were washed three time with ice cold PBS and detached from plates by incubation with ice cold PBS plus 10 mM EDTA. Cells were pelleted at 1,000 × g for 5 minutes at 4°C and cell pellets were flash frozen in liquid nitrogen and stored at −80°C. Cell pellets were lysed in 8M urea, 50mM Tris HCl, pH 8.0, 1X Roche Complete Protease Inhibitor and 1X Roche PhosStop. Lysates were quantified by Qubit fluorometry (Life Technologies). Samples were digestion overnight with trypsin. Briefly, sample reduction was done for 1h at RT in 12mM DTT followed by alkylation for 1h at RT in 15mM iodoacetamide. Trypsin was added at an enzyme:sample ratio of 1:20. Each sample was acidified in formic acid and subjected to SPE on an Empore SD C18 plate. For TMT labeling, after trypsin digestion ach sample was acidified in formic acid and subjected to SPE on an Empore SD C18 plate (3M catalogue# 6015 SD). Samples were dried out/lyophilized and reconstituted in 140 mM HEPES, pH 8.0, 30% acetonitrile. Fourty μL of acetonitrile was added to each TMT tag tube and mixed extensively. Tags were incubated at room temperature for 15 min. 30μL of label was added to each peptide sample and mixed aggressively. Samples were incubated in an Eppendorf Thermomixer at 300 rpm 25°C for 1.5h. Reactions were terminated by adding 8μL of fresh 5% hydroxylamine solution and 15min incubation at room temperature. Samples underwent high pH reverse phase fractionation following this conditions; Buffers: Buffer A: 10mM NaOH, pH 10.5, in water Buffer B: 10mM NaOH, pH 10.5, in acetonitrile. We used XBridge C18 colums, 2.1mm ID × 150mm length, 3.5μm particle size (Waters, part #186003023) hooked up to an Agilent 1100 HPLC system equipped with a 150μL sample loop operating at 0.3mL/min, detector set at 214 nm wavelength. Dried peptides were resolubilized in 150μL of Buffer A and injected manually. Fractions were collected every 30s from 1-49min (96 fractions total, 150μL/fraction). We analyzed by mass spectrometry 10% per pool for the full proteome in a nano LC/MS/MS with a Waters NanoAcquity HPLC system interfaced to a ThermoFisher Fusion Lumos mass spectrometer. Peptides were loaded on a trapping column and eluted over a 75μm analytical column at 350nL/min; both columns were packed with Luna C18 resin (Phenomenex). Each high pH RP fraction was separated over a 2h gradient (24h instrument time total). The mass spectrometer was operated in data-dependent mode, with MS and MS/MS performed in the Orbitrap at 60,000 FWHM resolution and 50,000 FWHM resolution, respectively. A 3s cycle time was employed for all steps. Data Processing Data were processed through the MaxQuant software v1.6.2.3. Data were searched using Andromeda with the following parameters: Enzyme: Trypsin Database: Uniprot Rat, Fixed modification: Carbamidomethyl (C) Variable modifications: Oxidation (M), Acetyl (Protein N-term), Phopho (STY; PO4 data only). Fragment Mass Tolerance: 20ppm. Pertinent MaxQuant settings were: Peptide FDR 0.01 Protein FDR 0.01 Min. peptide Length 7 Min. razor + unique peptides 1 Min. unique peptides 0 Second Peptides FALSE Match Between Runs FALSE The protein Groups.txt files were uploaded to Perseus v1.5.5.3 for data processing and analysis.

For phosphopeptide enrichment, we used two buffers Buffer A: 200μL of solution A + 800μL of acetonitrile /0.4% TFA and buffer B consisting of lactic Acid, 300 mg/mL in Buffer A. Dried peptides were resuspended in 100μL of buffer B. TiO2 tips (TiO2 enrichment kit, GL Sciences part # 5010-21312) were pre-washed/equilibrated with 20μL of buffer A, spun at 3,000 × g for 2 min, followed by 20 μL of buffer B, spin at 3,000 × g for 2min. Samples were loaded onto a TiO2 tip, spun at 500 × g for 2min. Flow through underwent loading onto a TiO2 tip, spun at 500 × g for 2min. Samples were washed with 20μL of buffer B, spun at 3,000 × g for 2min followed by 2 × 20μL of buffer A, spun at 3,000 × g for 2min. Peptides were eluted with 50μL of 5% NH4OH (pH 11.0) solution, followed by 50μL of 5% Pryrrolidine (pH 12.0). Eluates were frozen, dried in a lyophilizer and resuspended in 0.1% TFA for analysis.

### Stable isotope tracing experiments

250,000 cells per well were plated on a 6-well plate in 2 ml of DMEM (25 mM glucose, no pyruvate, penicillin/streptomycin, and 10% FBS). The next day, the cells were washed in 2ml of PBS and 2ml of DMEM media DMEM (25 mM glucose, no pyruvate, penicillin/streptomycin, and 10% dialyzed FBS) containing 1 mM ^13^C_3_ Pyruvate was added. After 12 hours, the medium was aspirated and 400 µl of lysis buffer (0.1M formic acid at 4:4:2 dilution (MeOH: ACN: Water)) was added. After 2 min on ice, 35µl of 15% NH_4_HCO_3_ was added and incubated for another 20 min on ice. Cells were then scraped, and lysates were transferred into pre-chilled 1.5ml centrifuge tubes, vortexed briefly, and spun at 15,060 rpm. for 30 min at 4°C. 390 µl of supernatant were then transferred into pre-chilled 1.5 ml centrifuge tubes and dried down using a Savant Speedvac Plus vacuum concentrator. Samples were resuspended in 120 µl of 4:1 (ACN: Water), sonicated for 5 minutes at 4°C, and centrifuged at 15,060 r.p.m. for 20 min at 4°C. 100 µl of supernatant was transferred into a pre-chilled LC-MS vial. 5 µl of this sample was injected on a HILIC-Z column (Agilent Technologies) and ^13^C labeled TCA intermediates were measured on an Agilent 6546 QTOF mass spectrometer that was coupled with an Agilent 1290 Infinity II UHPLC system (Agilent Technologies). The column temperature was maintained at 15 °C and the autosampler was at 4 °C. Mobile phase A: 20 mM Ammonium Acetate, pH = 9.3 with 5 µM Medronic acid, and mobile phase B: acetonitrile. The gradient run at a flow rate of 0.4 ml/min was: 0min: 90% B, 1min 90% B, 8 min: 78% B, 12 min: 60% B, 15 min: 10% B, 18 min: 10 %B, 19-23 min: 90% B. The MS data were collected in the negative mode within an m/z = 20-1100 at 1 spectrum/sec, Gas temperature: 225 °C, Drying Gas: 9 l/min, Nebulizer: 10 psi, Sheath gas temp: 375 °C, Sheath Gas flow: 12 l/min, VCap: 3000V, Nozzle voltage 500 V, Fragmentor: 100V, and Skimmer: 45V. Data were analyzed using Masshunter Qualitative Analysis 10 and Profinder 10 (Agilent Technologies).

### Bioinformatic Analyses

Gene ontology studies were performed with Cluego and Metascape. ClueGo v2.58 run on Cytoscape v3.8.2 (Bindea et al., 2009; Shannon et al., 2003). ClueGo was done querying GO CC and REACTOME, considering all evidence at a Medium Level of Network Specificity and selecting pathways with a Bonferroni corrected p value <10E-3. ClueGo was run with Go Term Fusion. To compare ontologies, we selected Metascape under multiple gene list and express analysis settings (Zhou et al., 2019). Analysis of microRNA targets was performed with the web tool MIENTURNET. High confidence network of targets was filtered using as parameters miRNA target interactions threshold at 2, FDR <0.06, cumulative weighted context++ score at 0.85, and probability of conserved targeting at 0.6 (Licursi et al., 2019).

### Statistical Analyses

Volcano plot p values were calculated using Qlucore Omics Explorer Version 3.6(33) without multiple corrections. Statistical analyses were performed with the engine https://www.estimationstats.com/#/ with a two-sided permutation t-test and alpha of 0.05 (Ho et al., 2019). Other statistical analyses were performed with Prism v9.5.0(525).

**Supplementary Figure 1.**
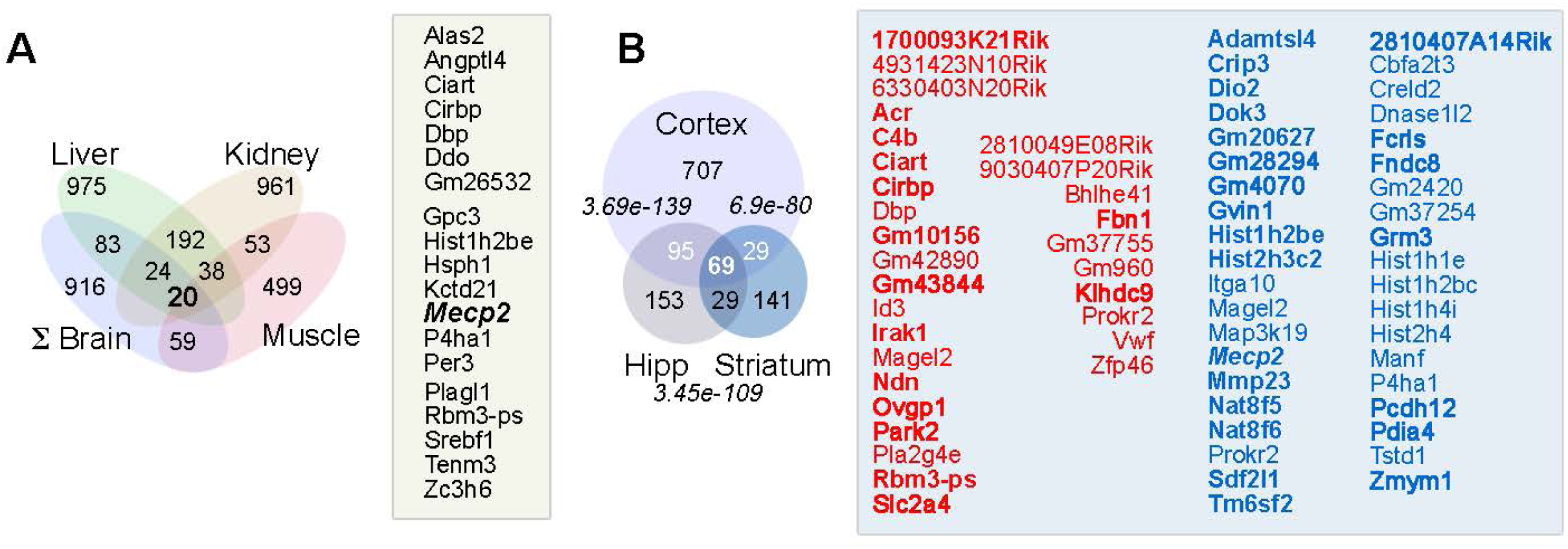
Overlap between *Mecp2^tm1.1Bird/y^* Tissue-Specific Coding Transcriptomes. **A.** Venn diagram of overlapping transcripts with altered steady-state levels common to all tissues. The 20 transcripts are listed in black font. **B**. Venn diagram of overlapping transcripts with altered steady-state levels in mutant tissue common to all brain regions. Hypergeometric p values for the different datasets overlaps. The 69 common transcripts are displayed in red and blue fonts depicting mRNAs whose steady-state levels are increased and decreased in mutants, respectively. Bold fonts depict transcripts also altered in Clemens et al. See Supplementary Table 1.

**Supplementary Figure 2.**
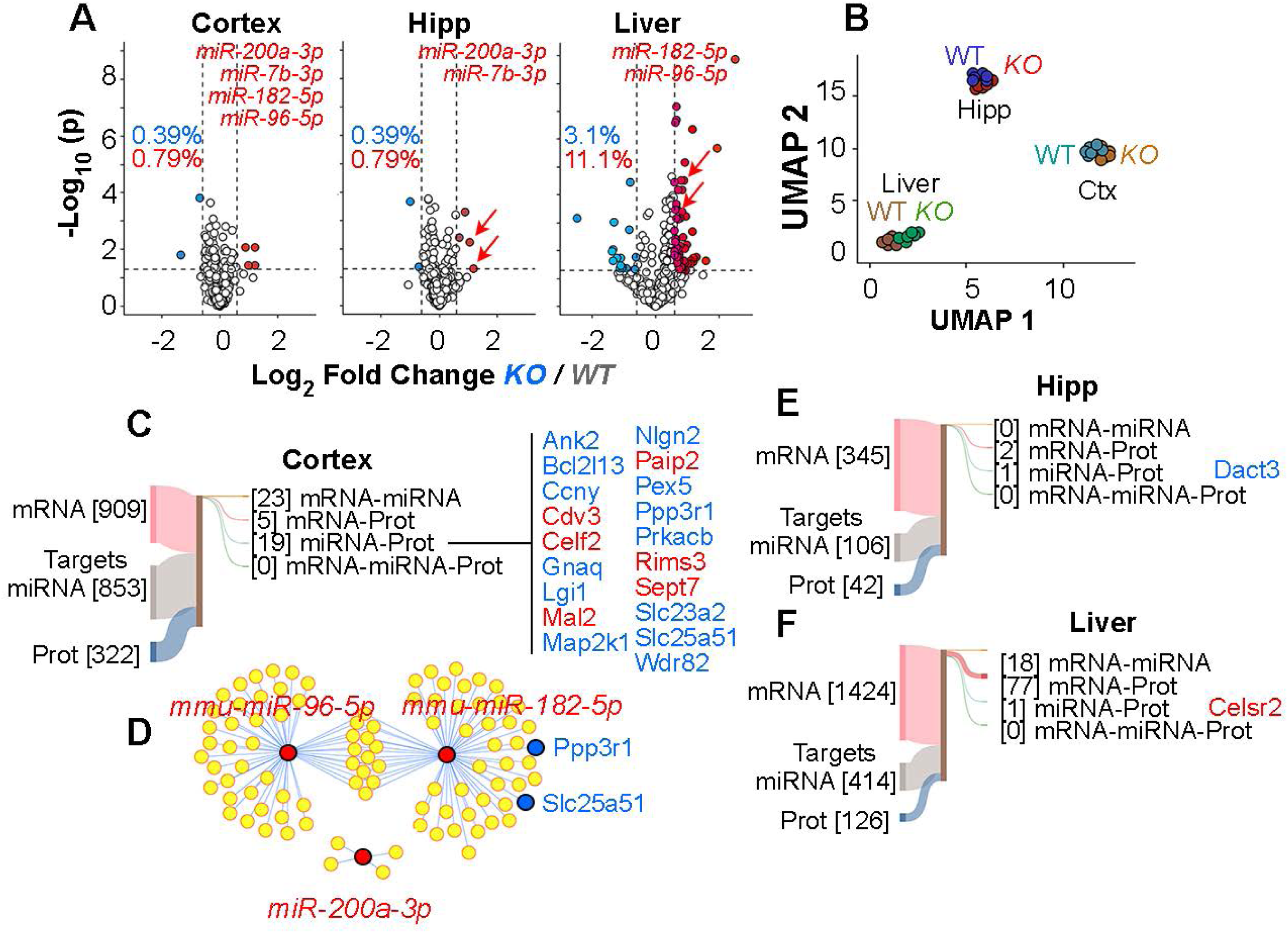
The *Mecp2^tm1.1Bird/y^* Tissue-Specific MicroRNA Transcriptomes. **A.** Volcano plots of microdissected cortex, hippocampus, and liver. Fold of change threshold is 1.5-fold and corrected p value cut-off is set at 0.05. Red and blue symbols represent transcripts whose steady-state levels are increased and decreased in mutants, respectively (n=6 animals of each genotype). **B**. UMAP analysis wild type and mutant tissue miRNA transcriptomes. **C, E** and **F**. Sankey graph depicting the overlap of predicted miRNA targets identified with the MIENTURNET tool (Licursi et al., 2019) and their intersection with the coding transcriptome and proteomes of the corresponding tissue. Predicted hits were not filtered by p values. Number in brackets represent the number of miRNA predicted hits intersecting with mRNAs or proteins altered in *Mecp2^tm1.1Bird/y^* mutant tissues. Blue font denotes transcripts/proteins with decrease steady-state levels, red font corresponds to transcripts/proteins with increased steady-state levels in mutant tissues. **D**. predicted cortex miRNA targets and their intersection with the cortex proteome using the MIENTURNET tool. Predicted hits were filtered by q values below 0.06. Blue dots are two proteins whose levels are decreased despite unaltered mRNA levels.

**Supplementary Figure 3.**
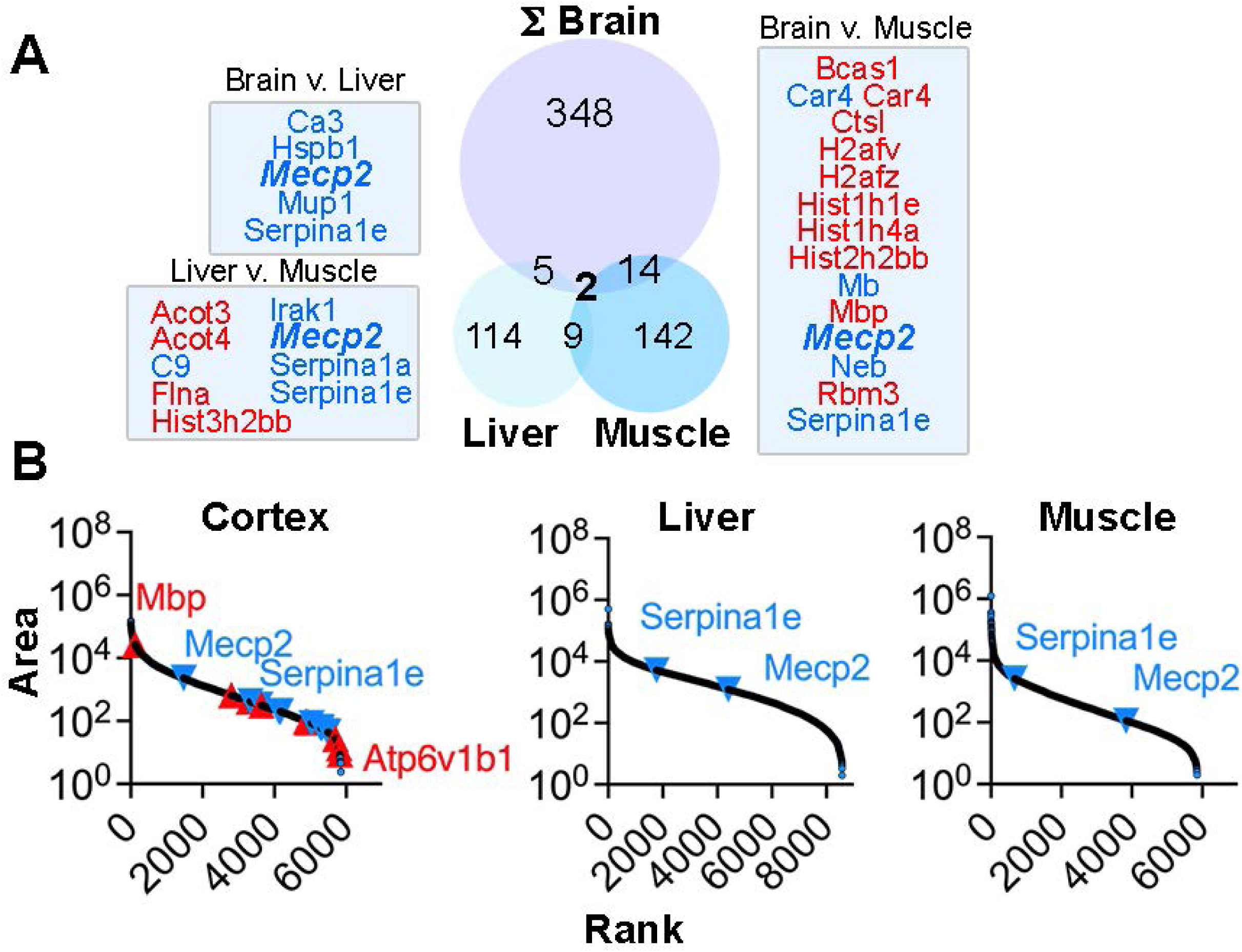
Overlap between *Mecp2^tm1.1Bird/y^* Tissue Proteomes. **A**. Venn diagrams of overlapping proteins with altered steady-state levels in mutant brain regions and the sum of all differentially expressed proteins in brain (ΣBrain) compared to muscle and liver mutant proteomes. Hypergeometric p values for the different datasets overlaps. Overlapping proteins are displayed in red and blue fonts depicting proteins whose steady-state levels are increased and decreased in each tissue. See Car4 increased in muscle and decreased in brain. See Supplementary Table 2. **B.** Protein level rank determined by mass spectrometry (area). The ten most increased (red triangles) and decreased (blue triangles) cortex proteins shown in Fig. 2B are depicted. Serpina1e, a protein decreased in all mutant tissues and Mecp2 are ranked in liver and muscle proteomes for comparizon.

**Supplementary Figure 4.**
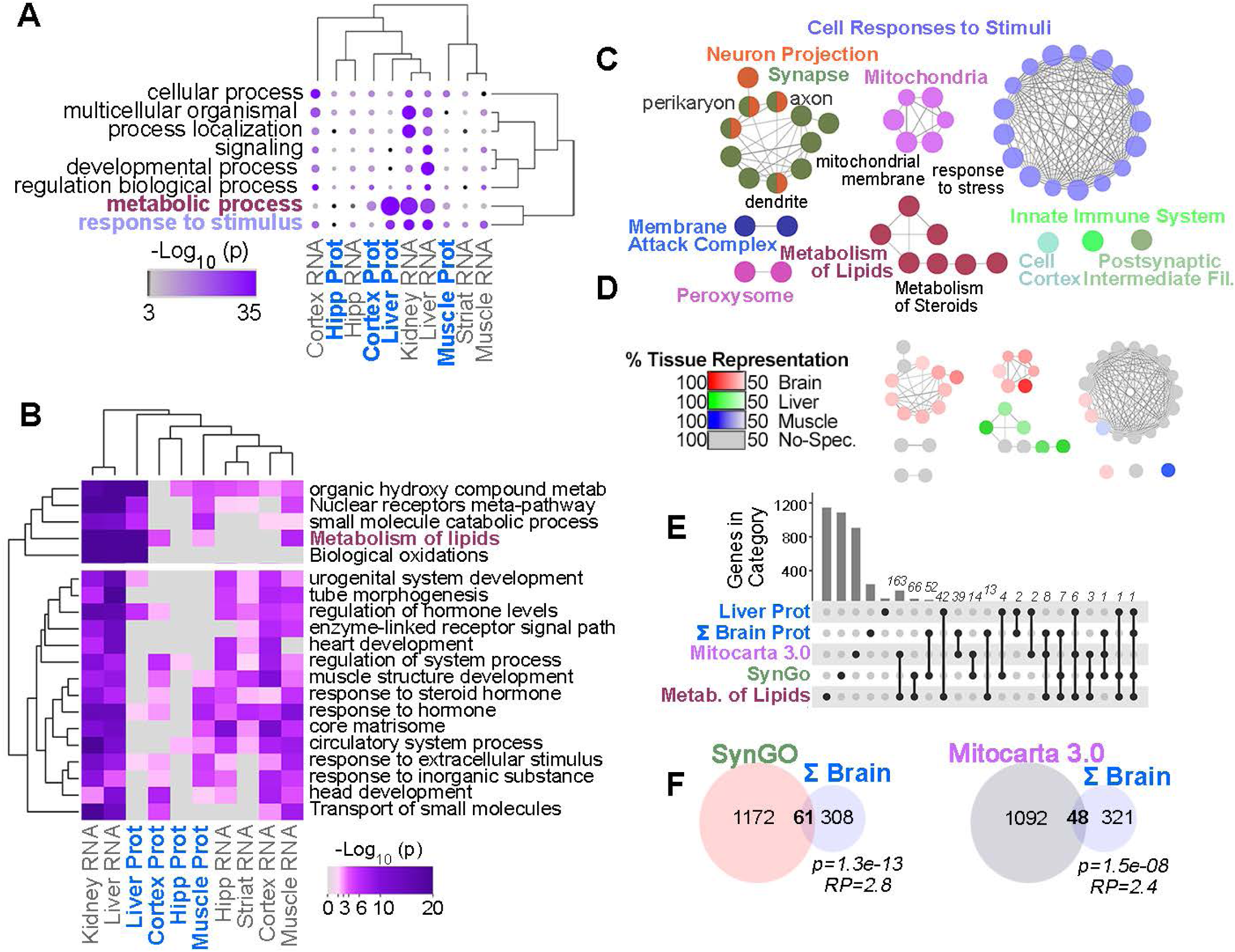
*Mecp2^tm1.1Bird/y^* Tissue Specific Proteomes and Transcriptome Converge on Metabolic and Mitochondrial Ontologies. **A.** Parental GO term ontologies represented in the differentially expressed proteomes and transcriptomes of mutant tissues. **B**. Ontologies, KEGG terms, and pathways enriched and shared by datasets used in A. For A and B accumulative hypergeometric p-values and enrichment factors were calculated and used for filtering. Significant terms were clustered into a tree using kappa-statistical similarities among their gene memberships. **C**. Interrogation of ontologies and pathways in mutant tissue proteomes using the ClueGO tool. All terms are significant with p<0.001 corrected with Bonferroni step down. **D**. Tissue representation of each ontology and pathway presented in C. Percentage represents % genes contributed by a tissue adjusted to input list size. **E**. UpsetR plot of overlaps between proteins annotated to SynGO, Mitocarta and metabolism of lipids (GO:0006629) present in liver and the sum of all proteins altered in mutant brain regions. Each black dot denotes the presence of unique overlapping hits shared by the categories in rows **F**. Venn diagram proteins with altered steady-state levels in mutant brain regions with the Mitocarta 3.0 and SynGO datasets. Hypergeometric p values for the datasets overlaps and the representation factor (RP. See Legend to Fig. S7B) indicating overlap above what is expected by chance. See Supplementary Table 3.

**Supplementary Figure 5.**
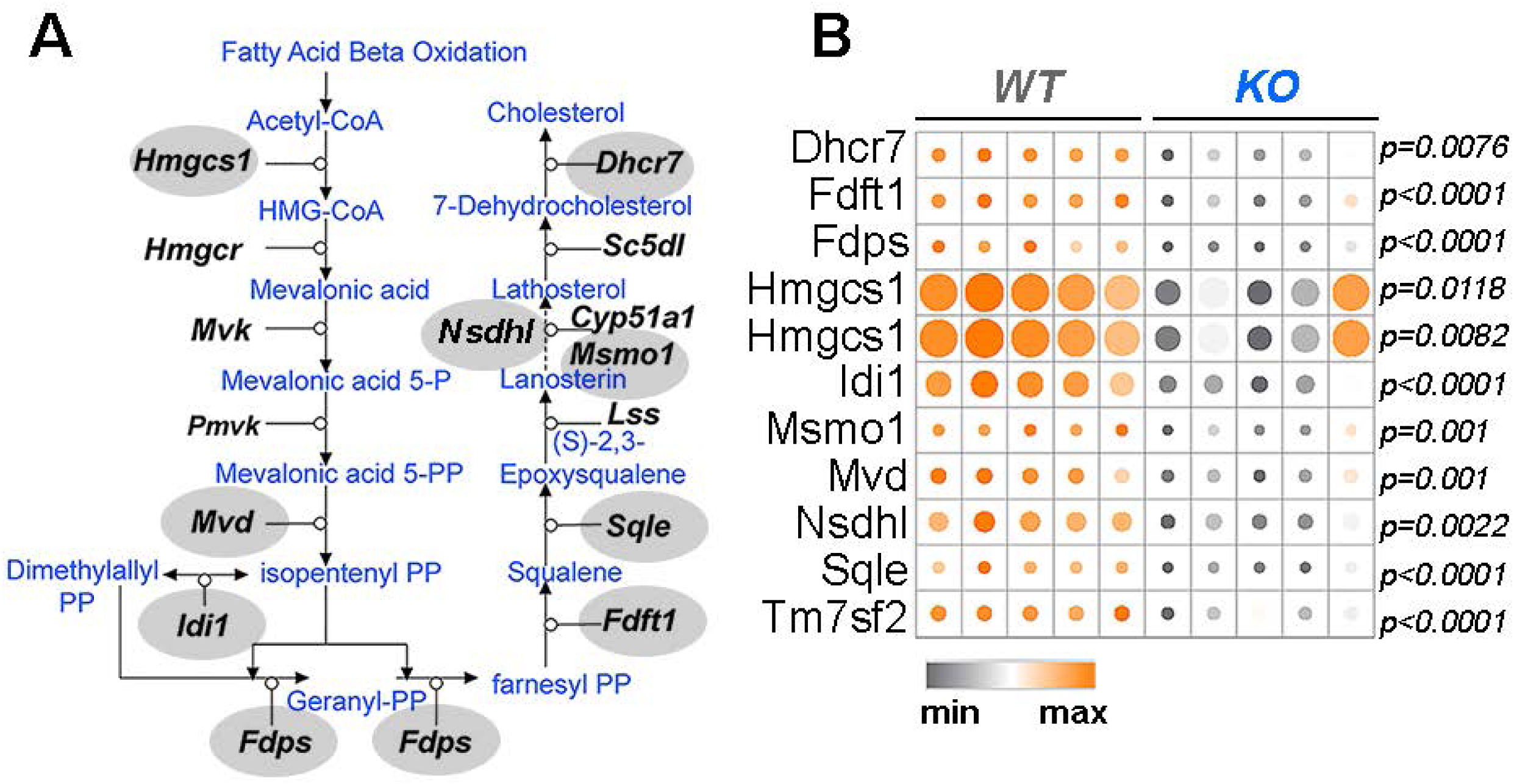
The *Mecp2^tm1.1Bird/y^* Liver Proteome Decreases the Steady-State of Cholesterol Synthesis Enzymes. **A.** Adapted cholesterol biosynthesis pathway (WP197). Enzymes decreased in mutant liver are marked by grey oval. **B**. heat map of the log_2_ normalized expression levels for enzymes depicted in A in wild type and *Mecp2^tm1.1Bird/y^* liver proteome. Each column is an animal. p values were obtained with two-sided permutation t-test with 5000 bootstrap samples.

**Supplementary Figure 6.**
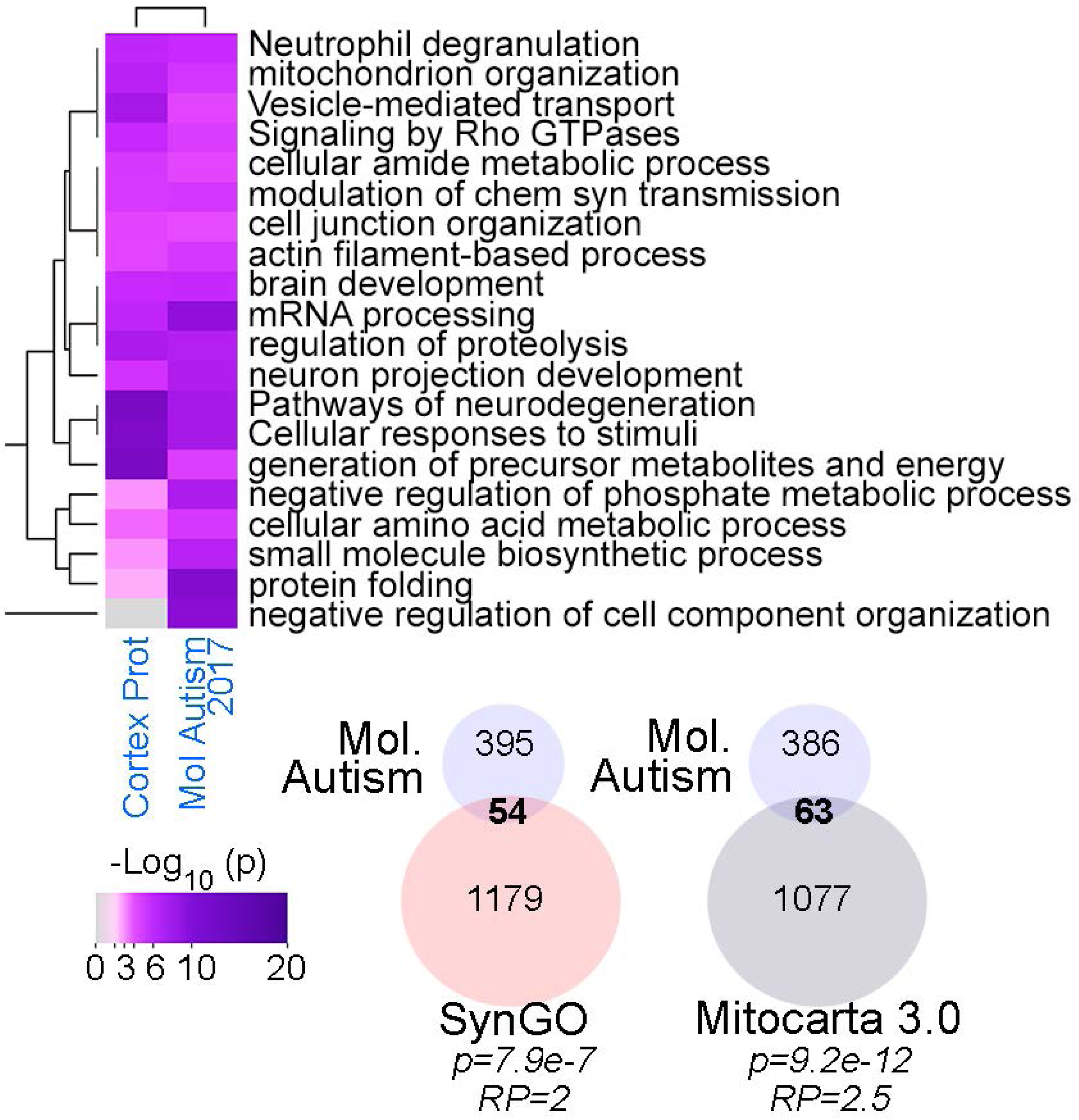
Ontological Convergence between *Mecp2^tm1.1Bird/y^* Microdissected Cortex and *Mecp2^Jae/y^* Isocortex Proteomes. Ontologies, KEGG terms, and pathways enriched and shared by cortex tissue datasets. Accumulative hypergeometric p-values and enrichment factors were calculated and used for filtering. Significant terms were clustered into a tree using kappa-statistical similarities among their gene memberships. Venn diagram proteins with altered steady-state levels in the *Mecp2^Jae/y^* isocortex proteome with the Mitocarta 3.0 and SynGO datasets (Koopmans et al., 2019; Rath et al., 2021). Hypergeometric p values for the datasets overlaps and the representation factor (RP, See Legend to Fig. S7B) indicating overlap above what is expected by chance.

**Supplementary Figure 7.**
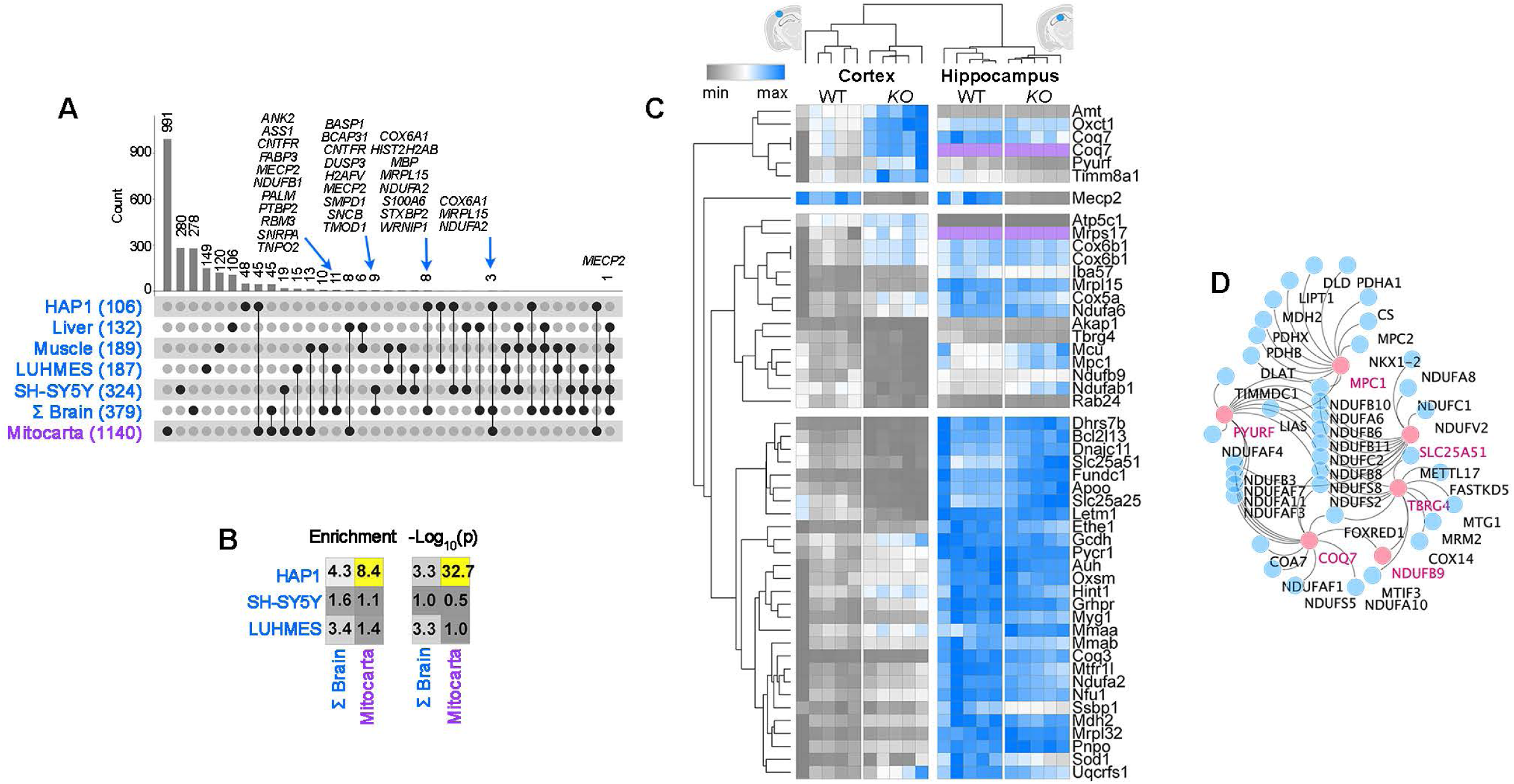
Mitochondrial Protein Commonalities Among *Mecp2^tm1.1Bird/y^* Brain and *MECP2* Null Cell Lines Proteomes. **A**. UpsetR plot of overlaps between proteins in different *MECP2*/*Mecp2* proteomes and proteins in Mitocarta 3.0. Each black dot denotes the presence of unique overlapping hits shared by the categories in rows. **B**. Overlap between human *MECP2* cell line proteome with the ΣBrain *Mecp2* proteome and the Mitocarta 3.0 dataset. Hypergeometric p values for the datasets overlaps and enrichment (representation factor) indicating overlap above what is expected by chance. Representation factor is defined as (# of genes in common between two groups/(# of genes in group 1*# of genes in group 2)/ total # proteins in either the human or mouse proteome)). A representation factor > 1 indicates more overlap than expected by chance between two independent groups. See Supplementary Table 3 and 4. **C.** Kendal Tau hierarchical clustering of the 61 Mitocarta 3.0 annotated proteins in mutant brain regions. Purple color denotes missing values. Log2 protein levels are expressed as a gray to blue scale ranging from the lowest to the highest log2 value per row. **D.** Fireworks co-essentiality network analysis of genetic relationships among mitochondrial proteins altered in *Mecp2^tm1.1Bird/y^* cortex. Nodes in pink represent most changed mutant cortex mitochondrial proteins in C.

**Supplementary Figure 8.**
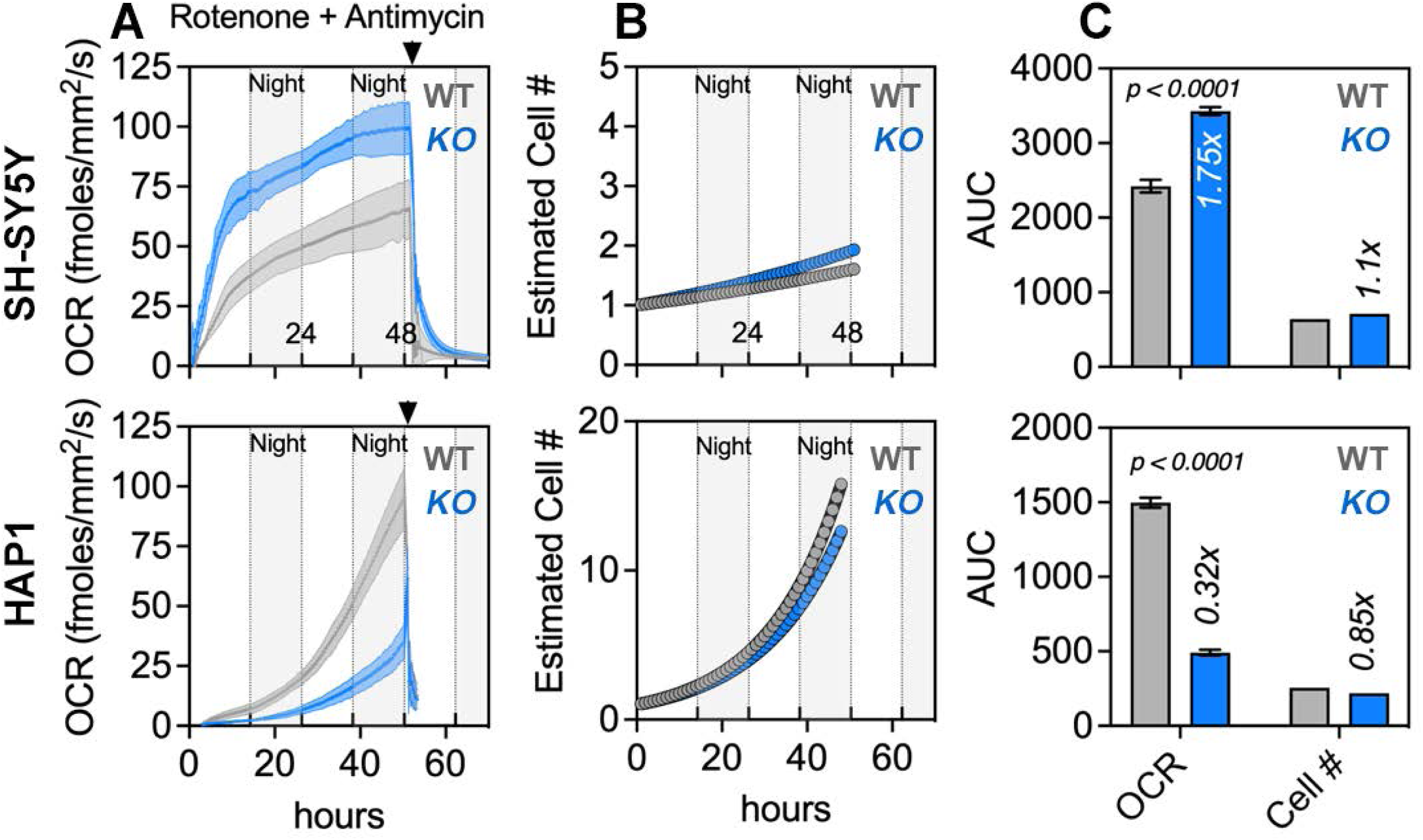
Long term oximetry of MECP2 null cells. **A**. Resipher oximetry determinations in wild type (gray) and *MECP2* mutant (blue) differentiated SHSY-5Y and HAP1 cells over a 48h period. Respiration was stopped by rotenone plus antimycin addition (arrowhead). **B**. Estimated cell count over the period of the oximetry based on the doubling time measured for each cell type and genotype. **C.** Total cell respiration and cell growth measured as the integrated area under the curve. Fold of change compared to wild type is presented as value(x). n=6-8, p value was calculated using t-test with Welch correction.

**Supplementary Figure 9.**
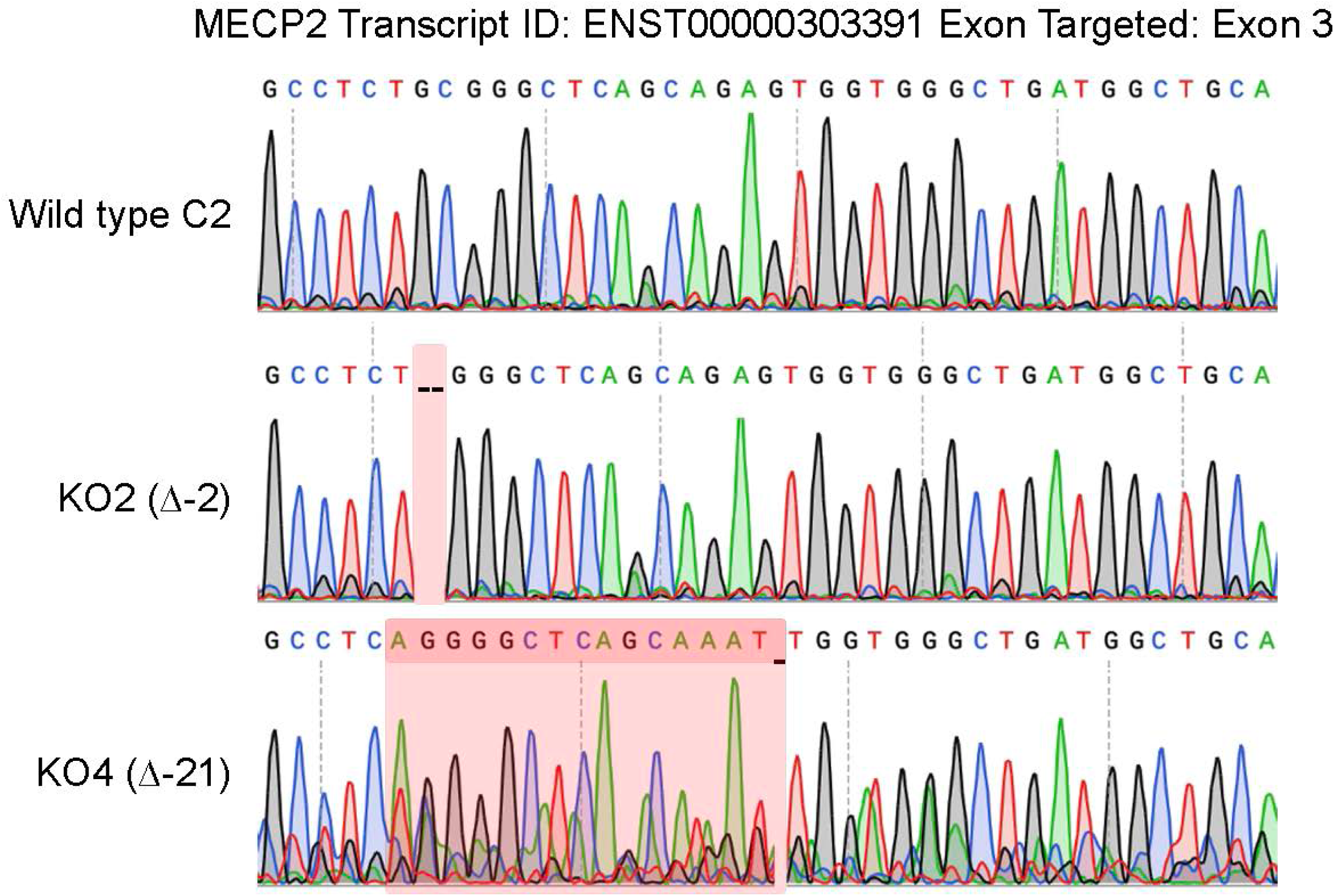
*MECP2* Gene Editing of SH-SY5Y Cells. CRISPR editing of the *MECP2* gene ENST00000303391. Pink shaded areas depict sequences that differ from the annotated *MECP2* gene. Null clone 2 has a deletion of 2bp and null clone 4 has a deletion of 21bp plus an insertion of 15bp. Blots for both *MECP2* mutant clones are presented in Figure 3A.

**Supplementary Table 1. *Mecp2^tm1.1Bird/y^* Tissue Transcriptomes.**

**Supplementary Table 2. *Mecp2^tm1.1Bird/y^* Tissue Proteomes.**

**Supplementary Table 3. Ontological Analyses.**

**Supplementary Table 4. *MECP2* Null Cell Lines Proteomes and Phosphoproteome.**

**Supplementary Table 5. Characteristics *MECP2* Null Cell Lines.**

## Acknowledgements

VF was funded by the Rett Syndrome Research Trust, the Loulou Foundation, and 1RF1AG060285. KSS holds a Postdoctoral Enrichment Program Award from the Burroughs Wellcome Fund and is supported by K00NS108539

